# The complexity dividend: when sophisticated inference matters

**DOI:** 10.1101/563346

**Authors:** Gaia Tavoni, Takahiro Doi, Chris Pizzica, Vijay Balasubramanian, Joshua I. Gold

## Abstract

Animals infer latent properties of the world from noisy and changing observations. Complex, probabilistic approaches to this challenge such as Bayesian inference are accurate but cognitively demanding, relying on extensive working memory and adaptive learning. Simple heuristics are easy to implement but may be less accurate. What is the appropriate balance between complexity and accuracy? We construct a hierarchy of strategies of variable complexity and find a power law of diminishing returns: increasing complexity gives progressively smaller gains in accuracy. The rate of diminishing returns depends systematically on the statistical uncertainty in the world, such that complex strategies do not provide substantial benefits over simple ones when uncertainty is too high or too low. In between, there is a complexity dividend. We translate these theoretical insights into specific predictions about how working memory and adaptivity should be modulated by uncertainty, and we corroborate these predictions in a psychophysical experiment.

## Introduction

Animals make sequences of sensory observations to arrive at judgements about current and future states of the world. In dynamic environments, this process is challenged by two primary forms of uncertainty (Heilbron & Meyniel, 2018; Yu & Dayan, 2005; Behrens et al., 2007): (1) noise, which obscures the useful information in signals; and (2) volatility, which is the tendency of the world to undergo change-points that reduce the relevance of the past to the future. In general, noise can be mitigated by remembering past experience and extracting average trends. In contrast, change-points require forgetting the pre-change-point history. Accordingly, we expect inference in dynamic environments to benefit from working memory for past experiences and adaptivity to environmental dynamics.

Models of inference, including those proposed to account for human and animal decision-making, can differ widely in form, accuracy, and complexity, leaving open basic questions about when and why each of these models is relevant to how the brain solves these problems (Rao, 2004; Bogacz et al., 2006; Gold & Shadlen, 2007; Fearnhead & Liu, 2007; Krugel et al., 2009; Shi & Griffiths, 2009; Brown & Steyvers, 2009; Nassar et al., 2010; Gigerenzer & Gaissmaier, 2011; Ossmy et al., 2013; Wilson et al., 2013; Legenstein & Maass, 2014; Glaze et al., 2015; Brody & Hanks, 2016; Veliz-Cuba et al., 2016; Glaze et al., 2018). The goal of the present study was to identify fundamental principles governing when particular cognitive operations are important to perform inferences that are both effective and efficient; that is, sufficiently accurate but also consistent with computational and information-gathering constraints that lead to bounded rationality (Gigerenzer & Gaissmaier, 2011; Gershman et al., 2015; Ortega & Braun, 2013). We reasoned that computational complexity in models of inference can represent a cognitive cost (e.g., in terms of the amount of working memory and the degree of adaptivity) that under some conditions might outweigh the benefits of potential gains in accuracy.

To test this idea, we constructed a hierarchical class of inference models that can be rated in terms of both accuracy and computational complexity. At the top of the hierarchy is Bayesian inference, which uses a probabilistic framework to combine both noise and volatility into a strategy that makes the most accurate inferences about current and future states of the world based on all previous observations (Adams & MacKay, 2007; Fearnhead & Liu, 2007; Wilson et al., 2013; Glaze et al., 2018; Griffiths et al., 2012; Radillo et al., 2017). This model provides a maximum-accuracy benchmark for our analyses, but it also can require virtually unlimited computational resources and thus provides a maximum-complexity benchmark as well. By deriving increasingly simple approximations to exact Bayesian inference, we then constructed two nested families of models corresponding to mental strategies that have progressively lower adaptivity and memory requirements (Fig. 1). Both accuracy and complexity decrease along the hierarchy.

**Figure 1:**
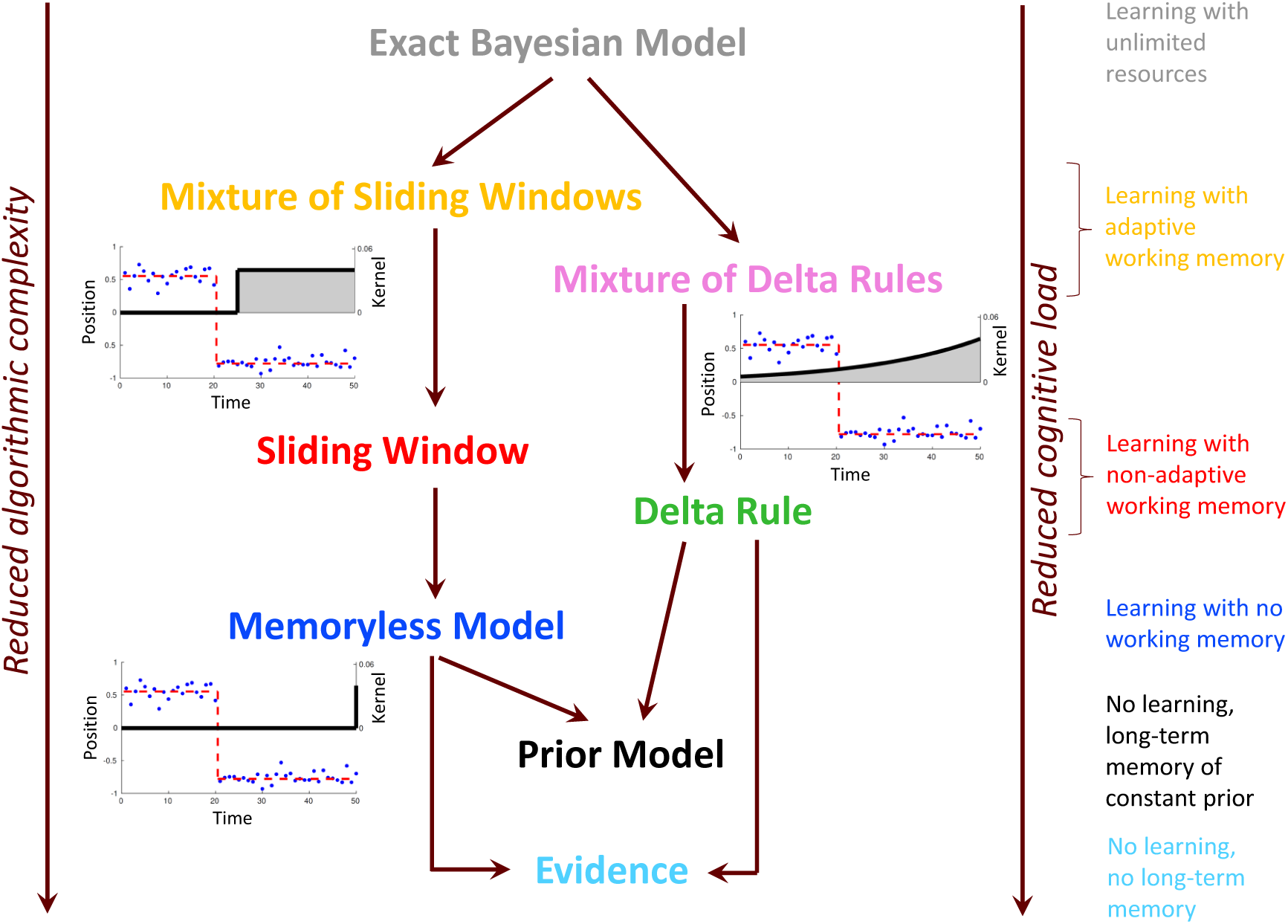
A hierarchy of cognitive functions maps to a hierarchy of inference strategies. Two nested families of inference strategies of decreasing algorithmic complexity can be derived from the exact Bayesian approach by progressively reducing requirements of memory and adaptivity (see also Fig. S1). We illustrate this in the context of inference from noisy observations (blue dots) of a latent variable *μ*_*t*_ (red dashed lines). The optimal Bayesian strategy balances prior belief against evidence integrated over temporal windows of all possible lengths, with each window weighted by the likelihood that the latent variable has been stable over that duration. The Mixture of Sliding Windows truncates the Bayesian model to a finite number of windows of fixed durations. The Delta Rules instead weigh past observations exponentially (examples of window and exponential integration kernels depicted as grey areas). The inferences from different Sliding Windows or Delta Rules are weighted optimally in the estimates of the mixture models. The Memoryless model simply combines current evidence with the prior and maintains no working memory (Dirac-delta kernel). The Prior model sticks to the prior belief regardless of evidence. The Evidence model follows the current evidence and ignores both prior beliefs and past evidence. The decrease in algorithmic complexity over this hierarchy of strategies mirrors a corresponding decrease in cognitive load (legend on the right-hand side).

We tested the performance of these nested models on tasks with varying noise and volatility and identified two fundamental principles. The first is a law of diminishing returns, whereby gains in accuracy become progressively smaller with increasing complexity, regardless of the amount of uncertainty in the environment. The second principle is a non-monotonic relationship between uncertainty and the complexity of the most efficient model: simple models are the most efficient when uncertainty is very high or low, whereas more complex models are useful at intermediate levels of uncertainty, when cues are both identifiable and helpful. Based on these principles, we generated specific, quantitative predictions about the most efficient use of adaptivity and working memory under particular uncertainty conditions. We then corroborated these predictions in an experiment probing human inferences under levels of noise and volatility that had not been explored previously. Overall, these results provide new insights into the cognitive processes that may be differentially engaged to perform efficient and effective inference under different conditions.

## Results

### A hierarchy of cognitive functions maps to a hierarchy of inference strategies

Numerous models have been proposed to solve inference problems in dynamic, noisy environments, ranging from complex, probabilistic ideal observers to simpler, heuristic update processes (Adams & MacKay, 2007; Nassar et al., 2010; Wilson et al., 2013; Sutton & Barto, 1998; Behrens et al., 2007). These models typically adapt to noise and volatility in their inputs by processing information over multiple timescales in a manner appropriate to the task conditions. Here we show that many of these models, and other plausible strategies, can be described parsimoniously in terms of two partially overlapping nested families that represent systematic simplifications of the Bayesian ideal observer (Fig. 1). The two families, which differ in terms of their working-memory demands, each include a progression from adaptive to non-adaptive, and less flexible, processing.

In general, a system for online inference aims to identify the current source of observations (the estimation problem) or to predict the next source (the prediction problem), in the presence of noise and unsignalled change-points, given all past and present data *x*_1:*t*_ = {*x*_1_,*…, x*_*t*_}. We consider a standard problem in which change-points in the source occur according to a Bernoulli process with a fixed probability *h* (volatility), and the source, characterized by a single number *μ*_*t*_ at a time *t*, generates observations with Gaussian variability (Fig. 2) (Wilson et al., 2013; Nassar et al., 2010). Noise in this generative process is measured by the ratio *R* between the standard deviation of the observations with respect to their source and the standard deviation of the sources across many change-points.

**Figure 2:**
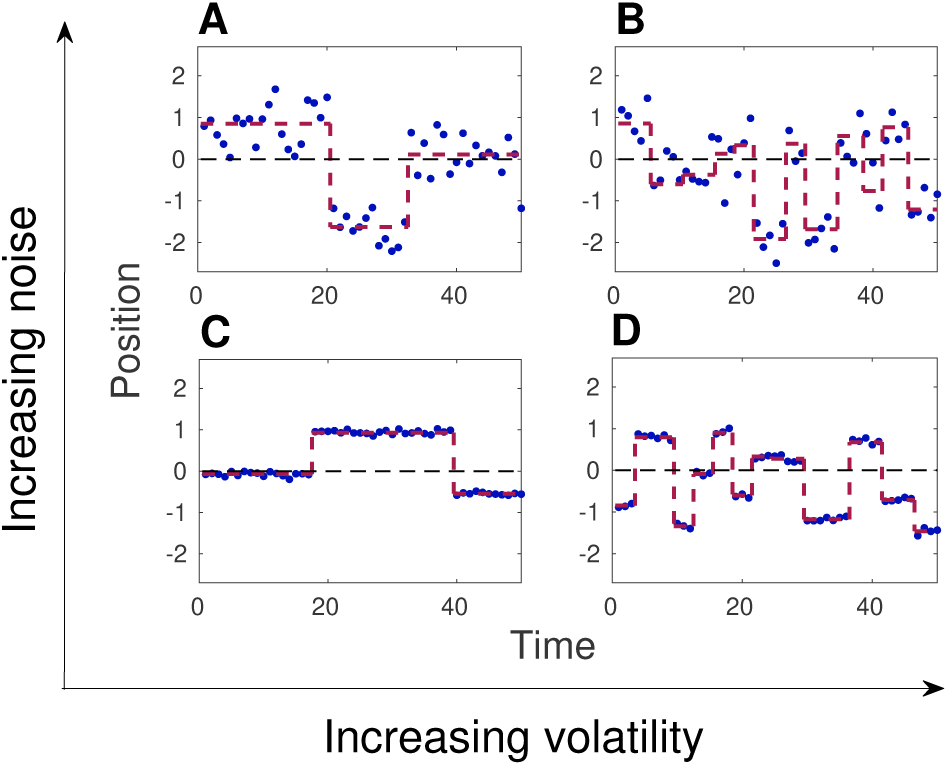
Gaussian change-point processes. Observations *x*_*t*_ (blue dots) are generated from a source positioned at *μ*_*t*_ (dashed red line) with Gaussian noise (SD = *σ*). The source is hidden to the observer and undergoes change-points at random times with probability *h* (volatility). At the change-points, *μ*_*t*_ is resampled from a Gaussian distribution centered at 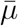 (dashed black line, stable over time) and with SD = *σ*_0_ = 1. Different panels show processes with different volatility (increasing from left to right) and noise *R* = *σ/σ*_0_ (increasing from bottom to top): **A**: *h* = 0.06, *R* = 0.45; **B**: *h* = 0.24, *R* = 0.45; **C**: *h* = 0.06, *R* = 0.05; **D**: *h* = 0.24, *R* = 0.05.

In this setting, the Bayesian ideal observer estimates the full distribution of the source in terms of two quantities: (1) the conditional probability *p*(*r*_*t*_|*x*_1:*t*_) of the run-length *r*_*t*_, which is the number of time steps elapsed at time *t* since the last inferred change-point in the source, and (2) the probability *p*(*μ*_*t*_|*r*_*t*_) that the source is *μ*_*t*_ given data observed over just the run-length *r*_*t*_. By multiplying these probabilities and summing over possible run-lengths, we can compute the probability that the source is *μ*_*t*_ given all the data:

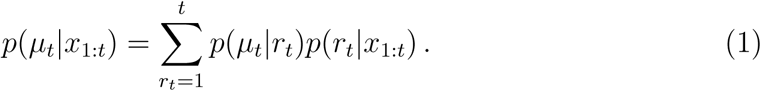

The Bayesian model computes *p*(*μ*_*t*_ | *r*_*t*_) and *p*(*r*_*t*_ | *x*_1:*t*_) exactly (Adams & MacKay, 2007; Fearnhead & Liu, 2007; Wilson et al., 2010). The optimal Bayesian estimate of the source, 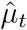, is then simply the expected value of *μ*_*t*_ in the conditional distribution (1). To optimally predict the next source, 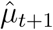, given this estimate, we must include the expected rate of change-points so that

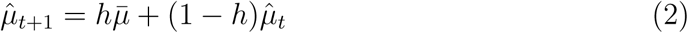

where 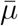 is the asymptotic average value of the source (Fig. 2). These Bayesian estimators minimize the mean squared error in both the estimation and prediction task.

The Bayesian model is computationally expensive: the time needed to make an estimate or a prediction grows linearly with *t*, because the model requires a sum over possible run-lengths (eq. 1; Wilson et al. (2013)). In cognitive terms, exact Bayesian inference requires working memory to increase with time. The computation is simplified by considering only a fixed set of run-lengths {*r*_1_,*…, r*_*N*_} chosen to minimize the mean squared error in the estimator. This reduction approximates the full Bayesian model with *N* computational units, each charged with generating an estimate of *μ*_*t*_ based on a sliding-window integration of past observations over the duration *r*_*i*_, combined with prior information on the average value of the source 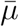 (eqs. 30, 31). This combination is chosen to guarantee that, when noise is high, the estimate is closer to the prior, which is more reliable than the sliding-window integration. Conversely, when noise is low, the estimate is primarily based on the sliding-window, which is more informative than the prior. Estimates computed by each unit are summed with a relative weight set adaptively by the posterior probabilities *p*(*r*_*i*_ |*x*_1:*t*_). As such, low/high volatility in the world will lead to preferential use of long/short sliding windows (Behrens et al., 2007; Fusi et al., 2007). This *Mixture of Sliding-Windows Model* is simpler than the full Bayesian procedure, but implementing it in the brain would still require extensive working memory, up to the longest run-length, and circuitry to compute the adaptive weights given to different run-lengths.

The working-memory load can be reduced by replacing the sliding windows with delta-rule updating units that weigh past observations according to an exponentially decaying kernel (eq. 33). Each delta-rule unit has a fixed time constant of decay linearly related to *r*_*i*_ that sets the timescale for information integration. The integration can be implemented recursively with a limited memory cost, through a simple update of the previous estimate by a fraction (learning rate) of the difference between the current observation and the previous estimate (eq. 32). The relative weight of the different delta rules in the combined estimate is again set adaptively by the posterior probabilities *p*(*r*_*i*_ |*x*_1:*t*_) (Wilson et al., 2013). This *Mixture of Delta-Rules Model* is simpler than the Bayesian model and requires less working memory than the Mixture of Sliding-Windows Model, but still requires computational resources to implement each of the delta-rules and to combine these with the correct adaptive weights.

The demand for computational resources in both model families is reduced dramatically by making them non-adaptive. This reduction amounts to considering a single *Sliding-Window* or a single *Delta-Rule*, each with a fixed timescale for integrating evidence. These models still require working memory to carry out the integration.

An even simpler inference strategy, which does not need working memory, estimates the source *μ*_*t*_ as a weighted average between the present observation *x*_*t*_ and the average source 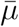 (eq. 34). This *Memoryless Model* is nested in the Sliding-Window Model, from which it is derived by choosing an evidence integration window of just one time-step. The Memoryless Model is the minimal model that learns and updates prior biases, or knowledge of stable features of the environment 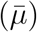, based on new evidence from rapidly changing variables (*x*_*t*_).

Both the Memoryless and Delta-Rule Models can be further reduced to the simple *Prior Model* 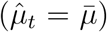 by setting the learning rate to zero. This Prior Model represents knowledge acquired, after a sufficiently long exposure to a given environment, about the constant or slow (stable across many change-points) features of the process generating the observations. Inferring and storing the slowly varying structure of the environment presumably still requires some cognitive effort and long-term memory resources.

Removing this last cognitive demand leads to a strategy that simply returns the current observation *x*_*t*_ as both an estimate of *μ*_*t*_ and a prediction of *μ*_*t*+1_. This strategy, which we call *Evidence*, can also be seen as the simplest possible model nested in both the Memoryless and the Delta Rule Models, because it is obtained from them by setting the learning rate to one.

These inference strategies form a hierarchy from the maximally accurate and cognitively demanding Bayesian model, to the maximally simple Evidence (Fig. 1). Each of these strategies estimates the source of observations and uses it to make predictions by computing a function that depends on (1) observations (*x*_1:*t*_), (2) fixed parameters of the environment (the average source 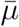, the volatility *h*, and the noise level *R*), and (3) model-dependent “meta-parameters” (the run-lengths and the learning rates). The simplifications giving rise to the two families of strategies from the full Bayesian model can be interpreted in terms of progressive reductions of cognitive demand.

### Adaptivity can be unnecessary when variability is low or high

When probability distributions are inferred from limited samples, complex models can generalize worse to new data than simple models (Balasubramanian (1997); Myung et al. (2000); Rissanen (1996, 1987); Barron & Cover (1991); Barron et al. (1998)). How to identify the model that best trades off fitting accuracy and generalization performance is the subject of a vast literature on model selection.

Here, we asked a different question. Even if a complex model has a lower prediction error, is the increase in complexity relative to a simple model “worth the effort”? One way to ask this question for nested model spaces like the hierarchy described in Fig. 1 is to find the best parameter configuration in the higher-dimensional parameter space of a more complex model, and to then determine whether the prediction error changes much if we vary the parameters to approach a simpler nested model. Formally, in an environment characterized by parameters {*e*_*l*_}, the best higher-dimensional model minimizes an error function *E* ({*e*_*l*_, *α*_*k*_}) with respect to its parameters {*α*_*k*_}. Thus, at the optimum 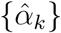, the gradient of the error with respect to the model parameters vanishes: 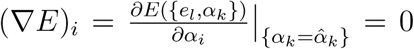. We can then characterize sensitivity of the error to the precise choice of parameters through the Hessian matrix

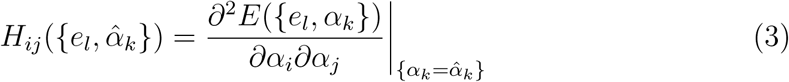

which evaluates the convexity of the error function at its minimum. The eigenvalues of *H* indicate how much the error *E* increases when moving away from the minimum in parameter space in the direction of the eigenvectors of *H*. Thus, a small eigenvalue indicates a combination of parameters that is substantially irrelevant for minimizing prediction error, whereas a large eigenvalue indicates a relevant combination of parameters (Gutenkunst et al., 2007).

We can characterize the irrelevance of the least-important parameter combination relative to the most-important one by evaluating the Redundancy, defined as the log ratio between the maximum and minimum eigenvalues of the Hessian matrix evaluated at the optimal parameters 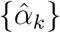:

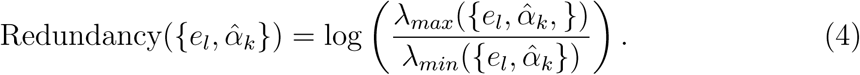

A Redundancy of *q* indicates that parameter deformations along the eigenvector associated to the least-relevant eigenvalue have a 10^*q*^-fold smaller effect on the error than deformations along the eigenvector associated to the most-relevant eigenvalue. If this irrelevant eigenvector points towards a simpler model nested within the parameter space, it suggests that the added complexity of the full model compared to the nested one is not necessary for good prediction performance. We can evaluate this Alignment as the (normalized) angle between the most irrelevant eigenvector and the most direct line from the optimal higher-dimensional model with parameters 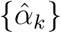 to the optimal lower-dimensional nested model (eq. 35, Methods).

We used this formalism to compare models attempting to predict the next value in the Gaussian change-point process in Fig. 2. The environmental parameters {*e*_*l*_} are the volatility *h* and the noise *R*, and *E* was chosen to be the mean-squared prediction error over 5000 time steps of the process for each choice of *h* and *R*. We computed the Redundancy of the mixture models with two Sliding Windows and two Delta Rules and their Alignment with respect to the optimal nested single Sliding Window and Delta Rule, respectively. Both mixture models have two parameters {*α*_1_, *α*_2_} describing effective learning rates, which are related to (1) the window length of evidence integration in the Sliding Windows (eqs. 30, 31), and (2) the timescale of the exponential evidence-integration kernel in the Delta Rules (eqs. 32, 33). The relative weight given to the two effective learning rates is determined adaptively at each time step based on accumulating experience. The non-adaptive single-unit models (one Sliding Window or Delta Rule) are obtained by setting *α*_1_ = *α*_2_. The Alignment, which takes values between 0 and 1, is simply the normalized angle *θ* between the most irrelevant parameter direction at the optimal mixture model and the line connecting the optimal mixture model to the optimal single-unit model (Fig. 3A).

**Figure 3:**
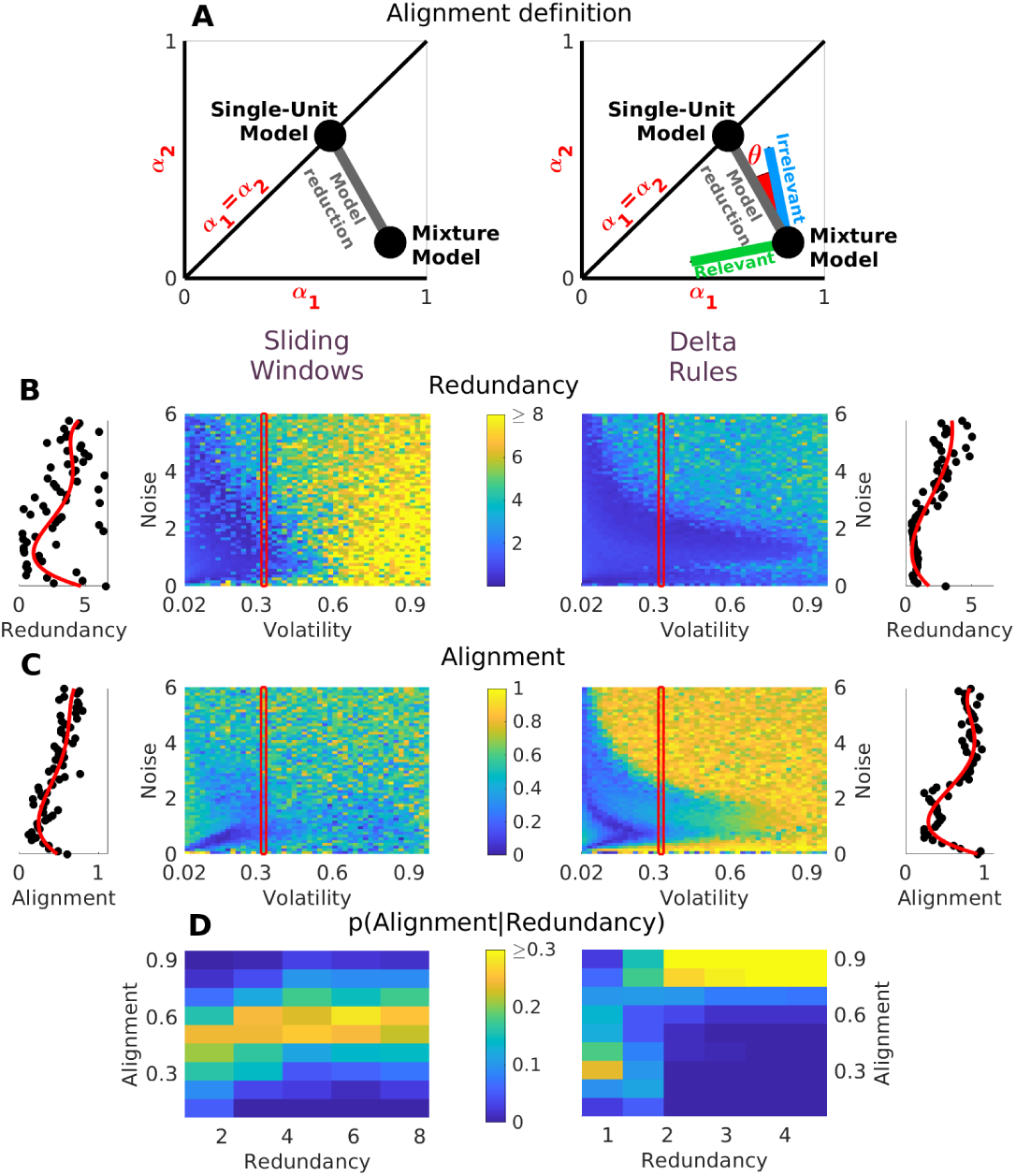
Adaptivity can be unnecessary when variability is low or high. **A:** Computation of the Alignment. (Left) Two-dimensional parameter space of the mixture models with two units defined by learning rates *α*_1_ and *α*_2_, and the embedded unidimensional space of the nested single-unit models (diagonal line *α*_1_ = *α*_2_). The optimal mixture model and optimal single-unit model (black dots) are indicated along with the parameter deformation leading from one to the other (gray line). (Right) Relevant and irrelevant parameter deformations that maximally or minimally change the prediction error moving away from the optimal adaptive mixture model. Alignment is defined as the normalized angle *θ* between the irrelevant deformation and the direction to the best non-adaptive single-unit model. **B:** Redundancy of the adaptive mixture models (left: Mixture of two Sliding Windows; right: Mixture of two Delta Rules) for a range of volatility and noise values in a change-point detection task (Fig. 2). Slices through the red inset windows are shown to the left and right, and show the non-monotonic trend of Redundancy with noise at low and intermediate volatility (red line: 4th-order polynomial fit): Redundancy is highest at low and high noise. **C:** Alignment of the irrelevant parameter deformation towards the non-adaptive nested single-unit model. Slices through the red inset windows are shown to the left and right and show a non-monotonic trend of Alignment with noise at low and intermediate volatility, similar to that observed for Redundancy. **D:** Probability distribution of Alignment values conditioned on Redundancy, sampled over tested volatility and noise values, shows that Alignment can be low at low Redundancy, but is typically higher at high Redundancy, indicating a rotation of the irrelevant parameter combination to align towards the non-adaptive nested model when Redundancy of the more complex model increases.

We found that the adaptive mixture models become redundant when noise and volatility are low so that inference is easy and complex strategies are unnecessary, and when noise or volatility are high so that inference is difficult, making complex strategies ineffective (Fig. 3B). When the models are redundant, the irrelevant parameter direction tends to align towards the simpler, non-adaptive single-unit models with optimally chosen parameters (Fig. 3C,D). This result suggests that when uncertainty is low or high simpler models will perform almost as well as complex ones. Sophisticated, adaptive inference matters only in a limited range of environmental conditions, characterized by relatively low volatility and intermediate noise (Fig. 3B,C).

### A power law of diminishing returns

The results in the previous section suggest that complex solutions to on-line inference problems may not always be worth the effort. To investigate this possibility quantitatively, we define the algorithmic complexity of an inference strategy in terms of the average number of computational operations required to implement it:

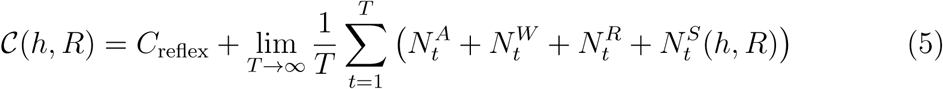

Here *T* is the total number of observations, 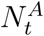 denotes the number of arithmetic operations and, 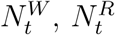, and 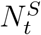 denote the number of memory-related operations (writing, reading, and storing, respectively) required to make a prediction at time *t*. Using conventional silicon hardware, multiplication typically requires more steps than addition, but here we will count elementary arithmetic operations as having unit algorithmic cost. We interpret the sum of these terms as an estimate of the reflective cost of making a decision, whereas *C*_reflex_ can be interpreted as a purely reflexive component that represents the irreducible cost of an action and is constant across models.

We evaluated the complexity of the different classes of models in Fig. 1 (Table 1 and Fig. 4A). For the non-parametric Bayesian Model, the reflective cost grows linearly with the number of observations because the entire past provides a probabilistic context for each prediction or estimate; thus the complexity 𝒞 diverges to infinity. The other models are parametric and have constant complexity that is partly related to the number of free parameters. This distinction is consistent with other measures of complexity like predictive information, which shows qualitatively different asymptotic behaviors for non-parametric versus parametric models (Bialek et al., 2001b,a). The complexity of models that involve mixtures of *N* Sliding Windows or Delta Rules grows quadratically in *N* (Table 1). Thus, the two-parameter adaptive mixture models (two Sliding Windows or Delta Rules as in Fig. 3) are more complex than the one-parameter non-adaptive models (Sliding Window, Delta Rule and Memoryless Models). These in turn are more complex than the two models with no parameters (Prior and Evidence). Models with the same number of parameters typically show smaller differences in complexity. These trends are qualitatively consistent with other notions of complexity from Bayesian model selection, information geometry, and data compression, for which the leading-order term of model complexity grows with the number of parameters, and lower-order terms depend on the model’s functional form (Schwarz, 1978; Balasubramanian, 1997; Myung et al., 2000; Rissanen, 1984, 1996; Barron & Cover, 1991; Barron et al., 1998). However, unlike those notions of complexity, algorithmic complexity can be applied readily to the kinds of deterministic models considered here.

**Table 1:** Asymptotic mean numbers of operations (*A*: arithmetic, *W* : memory writing, *R*: memory reading, *S*: memory storing) that determine the algorithmic complexity (eq. 5) of each model in the estimation problem. For the Mixture models, *N* denotes the number of units (set to 2 in this study) and *H*_2_ = *H*[*N* − 2] denotes the Heaviside step function centered in *N* = 2, which is equal to 1 for *N* ≥ 2 and to 0 for *N* = 1. Note that, for *N* = 1, the complexity of the Mixture models reduces to the complexity of the respective single-unit models. For the Sliding Windows, {*r*_*i*_(*h, R*)}, *i* = 1,…,*N*, is the set of run-lengths that minimizes the mean squared error of the model estimates given environmental volatility *h* and noise *R*.

| | $\langle N^A \rangle$ | $\langle N^W \rangle$ | $\langle N^R \rangle$ | $\langle N^S \rangle$ |
| --- | --- | --- | --- | --- |
| Evidence | 0 | 0 | 0 | 0 |
| Prior | 0 | 0 | 1 | 1 |
| Memoryless Model | 3 | 0 | 3 | 2 |
| Delta Rule | 3 | 1 | 3 | 2 |
| Sliding Window | 7 | 2 | 6 | $\lfloor r(h, R) \rfloor + 4$ |
| Mixture of $N$<br>Delta Rules | $3N + (7N^2 + 14N - 1)H_2$ | $N + (2N + 1)H_2$ | $3N + (6N^2 + 7N)H_2$ | $2N + (2N^2 + 2N + 1)H_2$ |
| Mixture of $N$<br>Sliding Windows | $7N + (7N^2 + 14N - 1)H_2$ | $N + 1 + (2N + 1)H_2$ | $6N + (6N^2 + 7N)H_2$ | $3N + \max_i (\lfloor r_i(h, R) \rfloor) + 1 + (2N^2 + 2N + 1)H_2$ |

**Figure 4:**
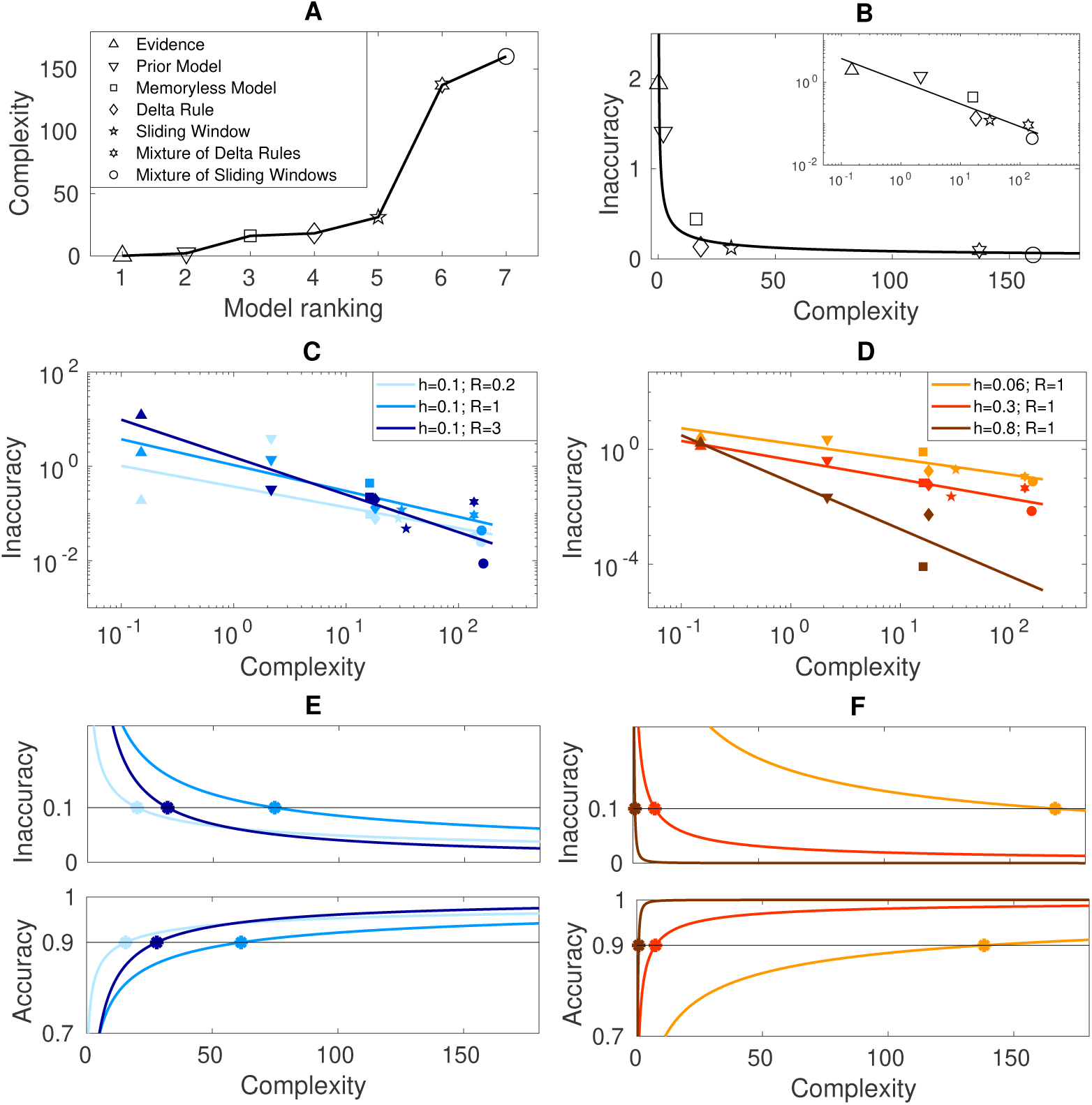
Diminishing returns from increasing complexity. Results are shown for the prediction task (inference of the next position of the source *μ*_*t*+1_). **A**: Algorithmic complexity (eq. 5) for models in Fig. 1. The mixture models take a weighted combination of evidence integrated over two timescales. The exact Bayesian Model has infinite complexity by our measure and is not shown. **B**: Inaccuracy (eq. 6) decreases as a power law in the Complexity (eq. 5), shown here for volatility and noise levels *h* = 0.1 and *R* = 1. Inset: linear fit on a log-log scale. See also Fig. S2 for goodness-of-fit statistics. The exponent in the power law varies with **C:** noise and **D:** volatility. **E**: Scaling of Inaccuracy and Accuracy (eq. 7) with Complexity for fixed volatility and varying noise. Color code and scaling exponents for each condition taken from panel (**C**). The convex/concave curves of Inaccuracy/Accuracy versus Complexity indicate a law of diminishing returns. Horizontal black lines indicate the threshold for performance within 10% of the Bayesian optimum. Intercept with the scaling curve for each task condition indicates the minimum model complexity required to reach the performance threshold. **F**: Same as panel (**E**) for fixed noise and varying volatility. Color code and scaling exponents taken from panel (**D**).

We measured performance of a model in terms of its Inaccuracy: the difference in mean squared error between the predictions of the model and those of the Bayesian ideal observer, normalized by the Bayesian benchmark, for each combination of volatility *h* and noise *R*,

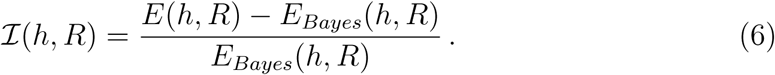

A vanishing Inaccuracy implies that the inference strategy performs as well as the Bayesian model. Across task conditions, we fit the Inaccuracy of the parametric models of Fig. 1 (with optimally chosen parameters) versus their Complexity. We found a power-law relationship between the two quantities (ℐ ∝ 1/𝒞^*b*(*h,R*)^) with an exponent that depends on volatility (*h*) and noise (*R*) in the underlying change-point process (Fig. 4B,C,D; goodness-of-fit summary statistics in Supplemental Fig. S2).

The power law for Inaccuracy as a function of Complexity implies a law of diminishing returns: increasing the complexity of a model gives progressively smaller improvements in prediction (flattening of Inaccuracy versus Complexity curves in the upper panels of Fig. 4E,F). To better visualize this effect, we also defined the Accuracy as the ratio

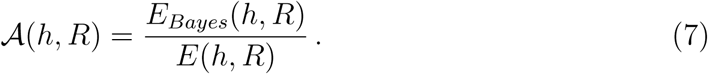

If the inference strategy performs as well as the Bayesian model the Accuracy will equal 1, whereas a model that makes very large errors on average compared to the Bayesian model will have an Accuracy tending to 0. The concavity of Accuracy as a function of Complexity (lower panels of Fig. 4E,F) again shows that prediction quality is a decelerating function of complexity.

In all task conditions, the curves (examples in Figs. 4E,F) show that prediction Accuracy is maximized and the Inaccuracy is minimized by using the most complex model. However, in the real world predictions typically do not have to be perfect – they just have to be good enough. We found that at both high and low noise (large and small *R*), low complexity models are already within 10% of the Bayesian optimum (light blue and dark blue lines in Fig. 4E). Likewise, when volatility is large, low complexity strategies perform almost as well as the full Bayesian model (red and brown lines in Fig. 4F). These results suggest that sophisticated inference procedures are only useful in a narrow range of conditions with an intermediate amount of noise and low underlying volatility. This conclusion is robust across a very wide range of thresholds for “good enough” performance, as seen by shifting the black threshold lines in Figs. 4E,F.

### Simple is usually best

The scaling laws identified in the previous section suggest that complex models are necessary only for a narrow range of conditions. To test this idea explicitly, we considered the nested hierarchy of models in Fig. 1 applied to the change-point detection task in Fig. 2. Over a wide range of volatility and noise levels, we selected the simplest of these models that achieved performance within 10% of the Bayesian optimum (Inaccuracy less than 0.1) in prediction and estimation tasks. Qualitatively similar results were obtained using alternative metrics and tolerance levels (Supplemental Figs. S3 and S4).

For prediction problems (Fig. 5A), extremely simple strategies reach nearly peak accuracy over a wide range of conditions. For example, when volatility is very high, the Prior model does nearly as well as the Bayesian predictor, because the world is so variable that past observations do not provide much useful information. When volatility is low, the underlying latent variables are persistent over time, so past observations become more useful for predicting the future. However, if noise is very high, observations are not reliable and so the Prior model is again nearly optimal. Conversely, if noise is very low observations are perfect, and so the present becomes fully predictive of the future without any need to consider the past. Thus, in the low-noise, low-volatility limit the Memoryless model (which simply balances current evidence with the prior) and the even simpler Evidence model (which simply follows the current observation and ignores the prior) are nearly as good as the Bayesian optimum. However, when volatility is low (so that there is something to learn from observations), and noise is intermediate (obscuring the latent variables, but not entirely), complex adaptive inference strategies are necessary for prediction performance that approaches the optimum. To summarize, when volatility is low there is a non-monotonic (“inverted-U”) pattern such that simple models are sufficient at low and high noise but complex strategies are needed at intermediate noise; when volatility is high simple strategies are always good enough.

**Figure 5:**
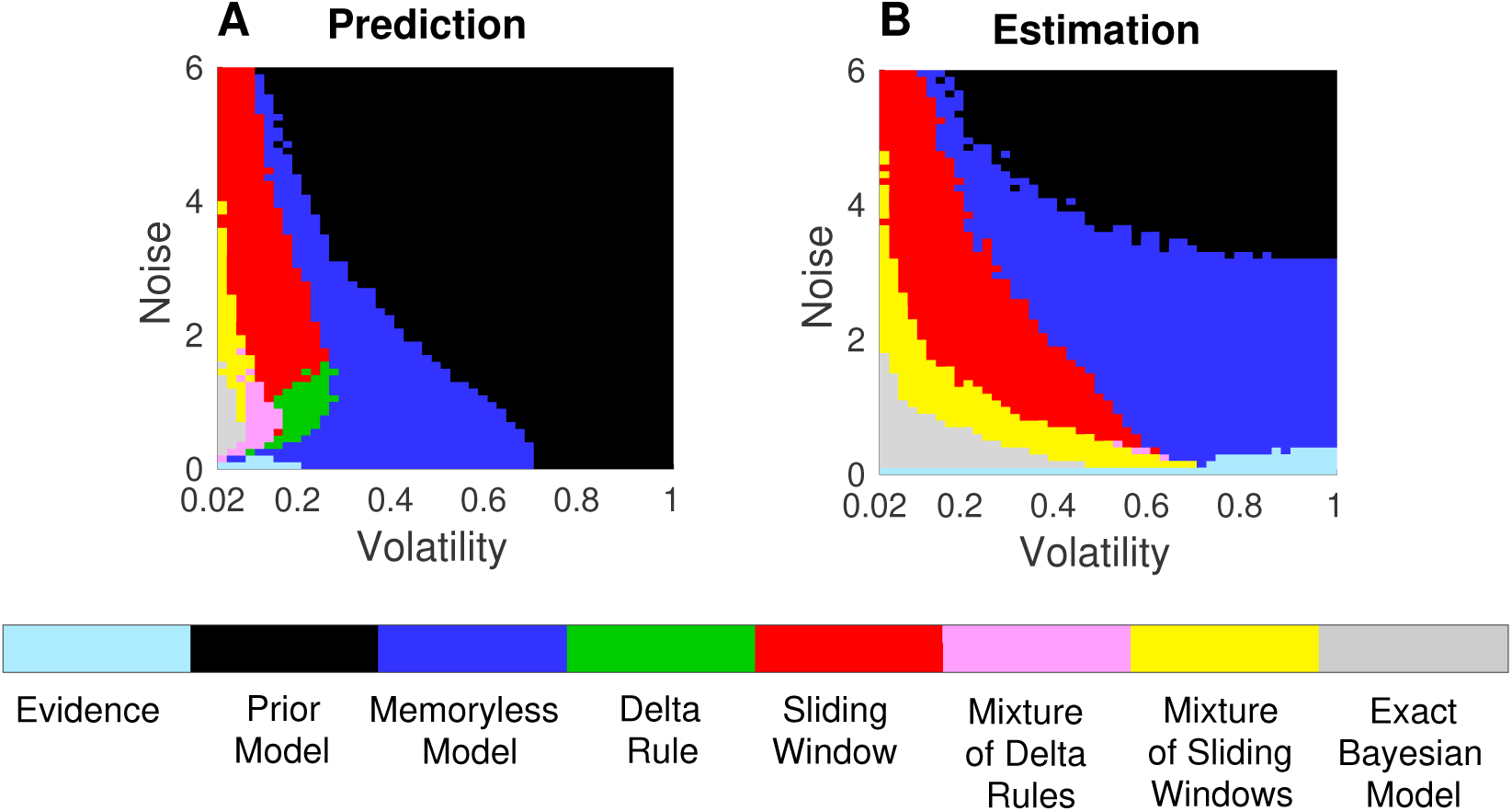
Simple inference strategies are usually sufficient. The color map shows the simplest strategy achieving performance within 10% of the Bayesian optimum (Inaccuracy < 0.1) for each combination of volatility and noise in the prediction (**A**) and estimation (**B**) tasks. Adaptivity and working memory are necessary in the gray, pink, and yellow areas; only working memory is required in the red and green regions. Extremely simple strategies (Evidence, Prior, and Memoryless Models) that use neither adaptivity nor working memory are sufficient in a vast domain of statistically easy and statistically difficult tasks. See also Figs. S3 and S4.

For estimation problems, slightly different patterns emerge (Fig. 5B). Like for prediction, simple strategies are almost as effective as the exact Bayesian model for high noise and high volatility. Meanwhile, when noise is very low the current sample always provides a good estimate of the current state, so the Evidence Model is effective regardless of volatility. As noise increases from zero at fixed volatility, complex models become useful to balance the current noisy evidence against past observations and the prior. But as noise becomes high (and observations unreliable), increasingly simple models are sufficient again to achieve near-optimal estimation performance. Thus, like for the prediction problem, when volatility is low there is an inverted-U relationship between the complexity required for good estimation and noise. However, comparing Fig. 5A (prediction) and Fig. 5B (estimation) we see that over much of the noise-volatility landscape estimation problems benefit more than prediction problems from the use of complex inference schemes.

### Optimizing cognitive engagement

Above, we selected the simplest model whose performance exceeded a hard threshold as compared to the optimal Bayesian strategy. It might be more realistic to imagine a smooth reward function that is low when the inaccuracy is high (ℐ ≫ 0) and high when the inaccuracy is low (ℐ → 0). This reward function can have a characteristic scale that sets the range of inaccuracies over which the animal receives a substantial reward. As a simple example, we can take the reward or performance level to be a Gaussian function of inaccuracy:

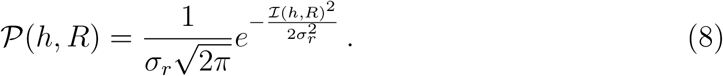

Thus, substantial rewards are obtained when ℐ is *O*(*σ*_*r*_) or smaller.

From Fig. 4B,C,D, we see that inaccuracy can be written as a power law in the complexity,

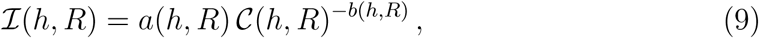

where *a* and *b* are fit parameters. Combining eqs. 8 and 9, then dividing by the complexity associated with a given level of inaccuracy from the fits, yields an expression for expected performance per unit complexity for each noise/volatility pair:

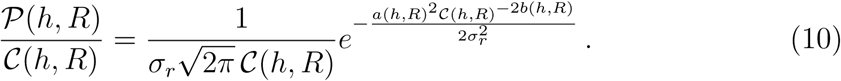

Because increased complexity in the inference strategy requires greater cognitive engagment, the ratio in (10) represents a trade-off between reward and cognitive cost per prediction or estimation. Because algorithmic complexity (eq. 5) can also be thought of as a qualitative estimate of the time required to make an inference, eq. 10 can also be interpreted as an estimate of the reward one can obtain per unit time (Shenhav et al., 2017; Vul et al., 2014; Schmidhuber, 2010; Gold & Shadlen, 2002).

The performance per unit cost can be optimized by maximizing the expression on the right hand side of eq. 10 with respect to the complexity *𝒞*. This procedure gives

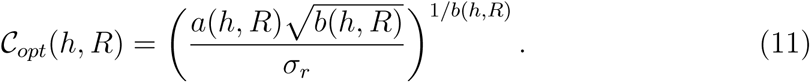

for the complexity, or, equivalently, cognitive cost, of the optimal inference strategy. Fig. 6 uses *a* and *b* measured from fits such as those in Fig. 4 to plot *C*_*opt*_ for prediction and estimation tasks across a range of volatilities and noise levels. The results confirm features seen in Fig. 5. For example, high complexity or cognitive engagement is needed only in a small subset of conditions, and follows an inverted-U trend with noise at low volatility (Fig. 6). Decreasing the width *σ*_*r*_ of the reward function decreases tolerance for large inaccuracies and thus broadens the domain where complex strategies are necessary. Changes in the reflexive component of complexity *𝒞*_reflex_ (eq. 5) leave the optimal reflective cost, and thus the optimal strategy, unaffected (Supplemental Information).

**Figure 6:**
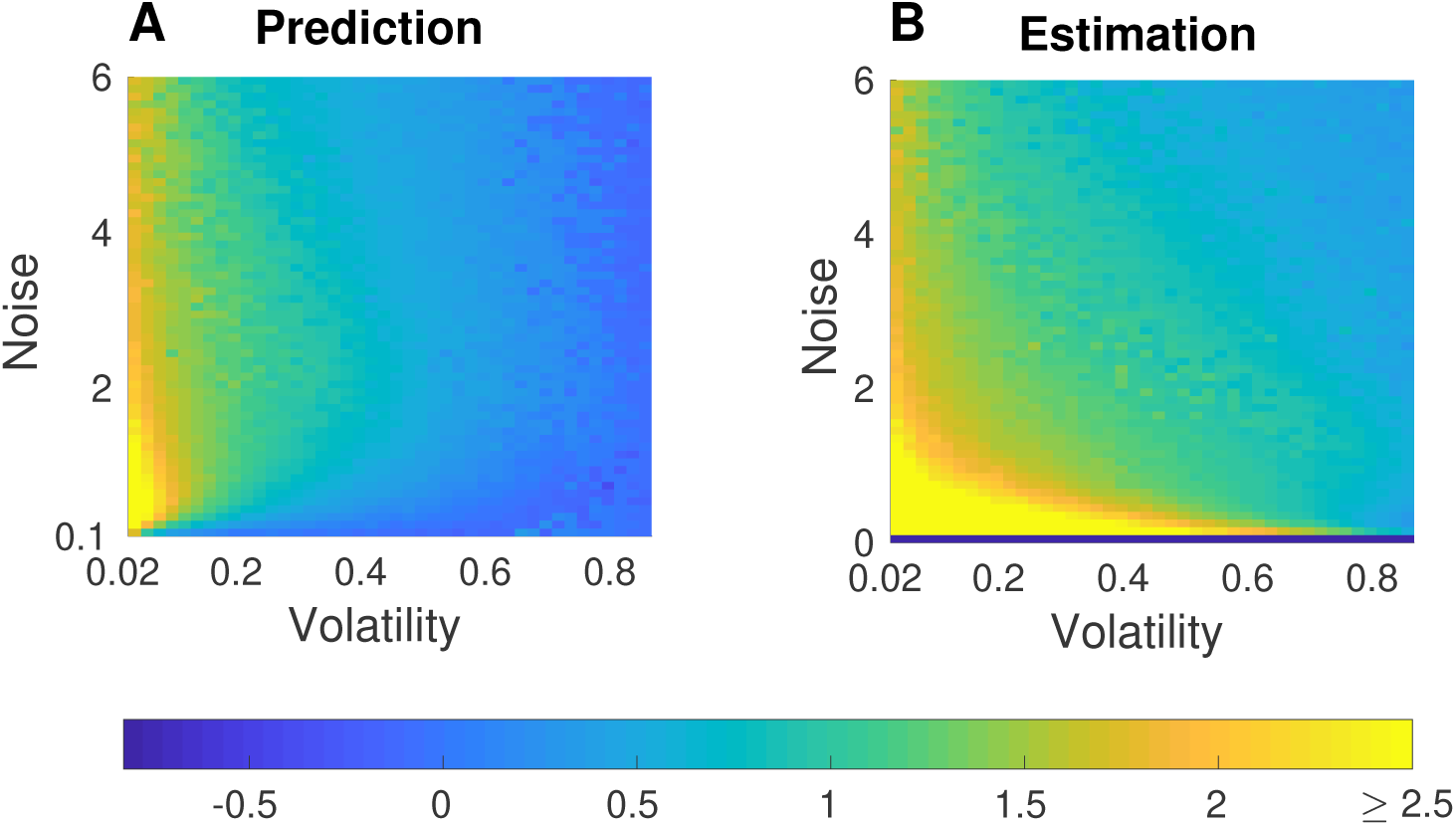
Optimal cognitive engagement. The colormaps show log_10_ *𝒞*_*opt*_ (eq. 11) as a function of volatility and noise, for the prediction (**A**) and the estimation (**B**) problems; *σ*_*r*_ = 0.1. High cognitive engagement is optimal only at low volatility and intermediate noise.

### Subjects modulate adaptivity and working-memory load as predicted by the theory

We used a psychophysical experiment to assess if and how our theory relates to human inference in terms of context-dependent uses of: (1) adaptivity, or the flexibility with which subjects change their integration time scale over past observations; and (2) working-memory load, or the maximum time scale over which subjects integrate past observations. Subjects were shown sequences of random numbers sampled from the kind of stochastic processes described in Fig. 2 and, on each trial, were asked to estimate the generative mean of the most recently observed number. Noise and volatility were held constant in blocks of trials and changed from block to block. One group of subjects was tested in three conditions of fixed (low) volatility and variable noise (circles in Fig. 7A). A separate group was tested in three conditions of fixed noise and variable volatility (diamonds in Fig. 7A). These six conditions substantially extend the range of volatility and noise probed in previous experiments, which focused on low-noise, low-volatility environments that require complex, adaptive inference (Wilson et al. (2013); Nassar et al. (2010); Krugel et al. (2009), small markers in Fig. 7A). From the individual responses in each condition, we estimated the integration kernel and integration time scale used by each subject to make an estimate of the generative mean at any given time-lag from the last change-point in the generative process. Adaptivity and working-memory load were then computed for each subject as the variance and maximum, respectively, of the integration time scales across different time-lags from a change-point. To best capture the relative, per-subject changes in these quantities across noise/volatility levels, adaptivity and working-memory load in any given condition were normalized by the maximum respective value across the three tested conditions. Theoretical values were computed in the same way as for the subjects, with simulated outputs from the most-efficient model (defined as the simplest model with ℐ < 0.1, as in Fig. 5B, but also using range of tolerances that might plausibly correspond to what human subjects would consider) in place of the subject responses.

**Figure 7:**
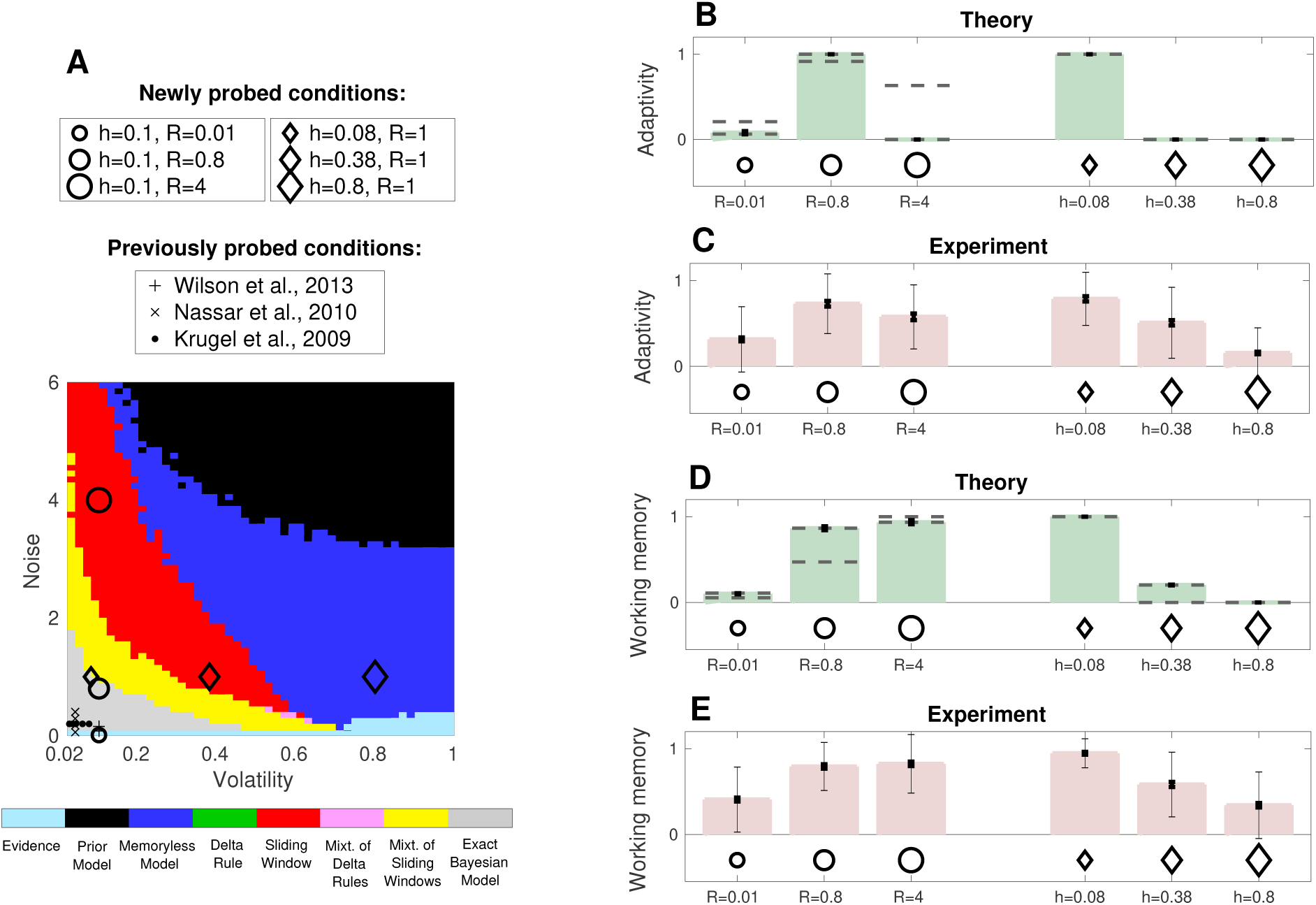
Subjects modulate adaptivity and working-memory load as predicted by the theory. **A**: Map of the volatility/noise conditions probed in this experiment (three conditions at fixed volatility *h* and variable noise *R*, and three conditions at fixed *R* and variable *h*) compared to conditions probed in previous experiments (see legend). Background colors indicate the theoretically most-efficient model (the simplest model with ℐ < 0.1) at each point of the volatility/noise space for the estimation task performed by the subjects (same as Fig. 5B). **B** and **C**: Mean normalized adaptivity ±SEM (small error bars) from the theoretically most-efficient model (**B**) and from 82/83 human subjects (**C**) performing the estimation task for each of the three noise (*R*)/volatility (*h*) conditions, respectively. Theoretical values were computed in the same way as for the subjects, with simulated outputs from the most-efficient model in place of the subject responses. For the colored bars, the most-efficient model was defined as the simplest model with ℐ < 0.1; dashed gray lines represent the range of values obtained using different tolerances (0.02 – 0.2; note the broad range for high noise). Thin error bars in C represent the standard deviation of the normalized adaptivity across subjects. Both the theory and data showed peak adaptivity at intermediate noise (left) and low volatility (right). **D** and **E**: Mean normalized working-memory load from theory (**D**) and 82/83 human subjects (**E**) performing the estimation task for each of the three noise/volatility conditions, respectively (plotted as in B and C). For both the theory and the data, the working-memory load is smaller at low versus intermediate and high noise and decreases with increasing volatility.

The subjects tended to adjust their use of both adaptivity and working memory across changes in noise or volatility in a manner that reflected key features of our theory (Fig. 7B–E). Specifically, we showed that, in principle, adaptivity is most useful for estimation tasks with intermediate noise and low volatility. Our subjects showed, on average, similar trends, with higher adaptivity for the intermediate versus low (one-tailed t-test, *p* < 10^−4^) or high (*p* = 0.003) noise and for the low versus intermediate or high volatility conditions (*p* < 10^− 4^ for both comparisons). Likewise, working memory is, in principle, most useful for estimation tasks with intermediate or higher noise and low volatility. Our subjects showed, on average, similar trends, with smaller working-memory loads at low versus intermediate and high noise conditions (*p* < 10^−4^ for both comparisons) and as a function of increasing volatility (*p*< 10^−4^ for all the comparisons). Differences across conditions were in general more pronounced for the theoretical than for the subject values. This slight discrepancy is attributable to two factors: first, we considered a finite set of models with discrete transitions between them, whereas subjects likely interpolate between different strategies, resulting in more variable differences in performance across conditions; second, the subjects’ responses were typically noisier than the simulated model outputs, resulting in more variable integration time scales. Even with these additional sources of variability in the subject data, our results show that our theoretical framework can be used to identify the task conditions in which different cognitive functions are most likely to be used by human subjects to solve inference problems.

## Discussion

### The models and the brain

We used a family of nested models and their mappings to particular cognitive functions to identify fundamental principles that govern the trade-off between the accuracy and simplicity of inference in noisy and changing environments. Each model we used can be seen as a particular implementation of a standard linear readout of the integrated activity in a population of neural units, which in the case of non-adaptive models reduces to a single neural unit (Wohrer & Machens, 2015; Shadlen et al., 1996; Haefner et al., 2013). The models differ in terms of the form of evidence integration they use and the degree to which they can adapt the time scale of this integration to the input. Both sets of properties have extensive and varied representations in the brain.

We considered exponentially decaying, sliding-window, and instantaneous (Dirac-delta function) integration kernels (implemented in the Delta-Rule, Sliding-Window and Memoryless Models, respectively). The exponentially decaying kernels correspond to the “*α*-synapses”, used widely in biophysical models of neuron spiking dynamics (Gerstner et al., 2014; Orhan, 2012). Implemented (with good approximation) as Delta Rules, they are also closely related to reward-prediction errors that are thought to be encoded by dopaminergic neurons and drive learning in the striatum and possibly elsewhere (Schultz et al., 1997; Schultz, 1998; Schultz & Dickinson, 2000; Waelti et al., 2001; O’Doherty et al., 2004; Behrens et al., 2007). This implementation, compared to exponentially decaying integration, has advantages in terms of working memory, because it effectively produces Markovian estimates of the source: each estimate depends only on the current observation and on the immediately previous estimate. The sliding-window kernels are more memory intensive, requiring representations of each sample used in the given window, or at least of the first and last samples in the window if implemented recursively. Such memory signals could, in principle, be based on persistent activity that maintains representations of a sequence of observations, such as those found in the prefrontal cortex network (Jacobsen & Nissen, 1936; Goldman-Rakic, 1995; González-Burgos et al., 2000; Kritzer & Goldman-Rakic, 1995; Funahashi et al., 1989; Arnsten et al., 2010). The Dirac-delta kernels can be implemented trivially without any working memory.

Adaptivity is achieved in our models using a bank of different integration timescales, consistent with multiple reports describing different integration timescales in the brain (Glascher & Büchel, 2005; Hasson et al., 2008; Bromberg-Martin et al., 2010; Bernacchia et al., 2011; Bornstein & Daw, 2012; Honey et al., 2012; Hasson et al., 2015; Meder et al., 2017; Scott et al., 2017; Runyan et al., 2017). In our formulation, the estimates obtained from these different integration timescales are weighted optimally and combined to produce a single output (Wilson et al., 2013, 2018). Consistent with this idea, learning rates with more relevance to an ongoing estimate of choice values have been shown to explain more variance in fMRI signals (Meder et al., 2017). This weighting process may be regulated by noradrenergic, cholinergic, and dopaminergic neuromodulatory systems, each of which has been linked to adaptive inference via pupillometry and other measures (Nassar et al., 2012; Krishnamurthy et al., 2016; Krugel et al., 2009; Aston-Jones & Cohen, 2005; Joshi et al., 2016).

An alternative hypothesis on how adaptive Bayesian inference might be approximated by the brain is based on particle filters and importance sampling (Fearnhead & Liu, 2007; Courville & Daw, 2008; Shi & Griffiths, 2009; Griffiths et al., 2012; Vul et al., 2014). In these approaches, a limited number of samples (particles) is used to represent the posterior distribution of the hidden state given the observations. Unlike in our models, in these approaches the hypothesis space for the hidden state varies in time, as new hypotheses are continuously sampled from their Bayesian posterior distribution given the observations. By contrast, in our Mixture Models the hypothesis space (set of run-lengths or integration timescales) is fixed in a given environment and adaptivity is achieved by assessing the different hypotheses differently in a time-dependent manner, based on the accumulated evidence. It would be useful for future work to compare the computational complexity of these different kinds of approaches to adaptive inference, which could help constrain our understanding of if and when they could be used in the brain.

Overall, our study provides a unified view of several plausible models of on-line statistical inference, showing that they can be regarded as special cases of a single formalism. This novel interpretation suggests a hierarchical (nested) organization of cognitive processes and a natural, efficient way in which the brain could engage or disengage them. Specifically, this organization implies that the brain could meet the demands of a wide range of different environments and tasks, by adjusting the parameters of a single, flexible inference process.

### Inaccuracy versus complexity trade-off

Each of our models is characterized by its complexity and by its inaccuracy compared to the exact Bayesian Model. By analyzing two nested families of models, we identified a power-law scaling of inaccuracy with complexity: ℐ ∝ 1/𝒞^*b*^. This scaling, with an exponent that depends on noise and volatility in the environment, implies a law of diminishing returns such that increasing the complexity of the inference strategy gives progressively smaller returns in prediction/estimation accuracy. This law is reminiscent of a similar result in rate-distortion theory: the minimum achievable distortion 𝒟 of a transmitted signal is a continuous, monotonically decreasing, convex function of the information transmission rate ℛ (Cover & Thomas, 2012). This universal property of rate-distortion functions implies that, independently of the source of information, increasing the communication rate confers diminishing returns in reconstruction accuracy at the receiver. In simple contexts, the rate is measured in bits as the mutual information, 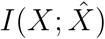, between the input *X* and output 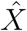of an information channel (Cover & Thomas, 2012). Noting that the distortion for a Gaussian channel (similar to our Gaussian source) scales as log *D* ∼ − ℛ (for distortions smaller than the variance of the samples) and that our inaccuracy scales as log ℐ ∼ − log 𝒞 suggests an interpretation of the log algorithmic complexity of our models as an effective transmission rate of information about the environment to a decision making “receiver”, who gathers this information to make inferences about the world. In this analogy, constraints on the inference algorithm, imposed by bounded rationality (Gershman et al., 2015), create a sort of information bottleneck (Tishby et al., 2000). The connection with information theory may provide new practical tools to help understand the diversity of strategies used across tasks and individuals to solve inference problems (Glaze et al., 2018). Such tools also have the potential to deepen our understanding of the diversity of deep neural networks where a power-law scaling between accuracy and computational complexity reminiscent of our findings has recently been identified (Canziani et al., 2016).

### Inverted-U relationship between cognitive demand and task difficulty

A key finding of our theory is that complex strategies that use adaptive processes and/or working-memory are necessary only for a restricted range of conditions characterized by low volatility and moderate noise, with working memory being useful across a slightly wider set of uncertainty levels, particularly towards higher noise. We showed that, under certain conditions, human inference also follows these basic trends: (1) adaptivity is reduced at low and high versus intermediate noise (when volatility is low), and decreases with increasing volatility; (2) working-memory is lowest at low noise and decreases with increasing volatility. Both the theory and the experiments highlight the fact that simple strategies are good enough when inference is easy, such as when the current evidence from the environment is highly reliable and thus historical information is not needed, or when inference is hard, such as when incoming information is so noisy or volatile that there is little information to gain from complex reasoning.

This “inverted-U” relationship between cognitive demand and task difficulty is reminiscent of a similar relationship between cognitive abilitites, like learning, and arousal state (Yerkes & Dodson, 1908; Phillips et al., 2004; Durstewitz & Seamans, 2008; Cools & D’Esposito, 2011; Arnsten et al., 2012). Several lines of evidence suggest that this relationship reflects the effects of neuromodulators like norepinephrine and dopamine on neural activity in the prefrontal cortex and perhaps elsewhere in the brain (Aston-Jones et al., 1999; Aston-Jones & Cohen, 2005; Arnsten et al., 2012). It is tempting to think that the statistical difficulty of a task might modulate activity in these brain areas similarly to arousal states, to engage or disengage mental resources in a way that best meets task demands.

An inverted-U relationship is also found in combinatorial optimization problems, suggesting that it might be a much more general phenomenon: NP-complete problems such as K-satisfiability, graph coloring, the traveling salesman, and the Hamiltonian path problem, have characteristic easy-hard-easy patterns in the computational complexity required to find a solution. Hard problems are typically clustered around a critical intermediate value of an order parameter, which marks a phase transition from solvability to unsolvability (Cheeseman et al., 1991; Mitchell et al., 1992; Hogg et al., 1996; Gent & Walsh, 1996; Hayes, 1997; Monasson et al., 1999; Cocco & Monasson, 2001; Biroli et al., 2002; Zdeborová, 2009). In a broad sense, this order parameter plays a role similar to environmental uncertainty in our inference task.

### Comparison with previous experiments and perspective

To the best of our knowledge, our experiment is the first to probe human inference for a wide range of noise and volatility conditions and to show how adaptivity and working memory are modulated by these widely different levels of uncertainty. The conditions tested in previous studies were restricted to low volatility and moderate noise (Fig. 7A). In line with our theory, those studies showed that human inferences were consistent with relatively complex, adaptive strategies (Wilson et al., 2013; Nassar et al., 2010; Krugel et al., 2009).

Specifically, in Wilson et al. (2013, 2018), human subject predictions were consistent with a Mixture of Delta-Rules strategy with two computational units in a Gaussian change-point task similar to the one considered here, when volatility was ∼ 0.1 and noise was ∼ 0.1. This model provided a better fit than a model with either one or three units. Note that the (*h* ∼ 0.1, *R* ∼ 0.1) point in the volatility-noise plane falls in the small region where the 2-Delta-Rule Model is the most efficient strategy for prediction tasks according to our theory, assuming plausible tolerances to errors for human subjects (between 2% and 10%); see Fig. 5 and Fig. S3. Likewise, in Nassar et al. (2010), a Delta-Rule Model with an adaptive learning rate fit human behavior on a similar Gaussian change-point task better than a Delta-Rule with a fixed learning rate, when volatility was ∼ 0.04 and noise ranged between ∼ 0.06 and ∼ 0.4. This result is in agreement with the adaptive domain in our map of efficient models for both prediction and estimation tasks, assuming a tolerance to errors in the same plausible [2% 10%] range (Fig. 7A). In Krugel et al. (2009), adaptive changes in learning rates were detected, in a probabilistic object-reversal task, for almost all tested subjects. In this task, the probabilities of object reversal (analogous to volatility) ranged between ∼ 0.008 and ∼ 0.08, and the fraction of trials in which the statistically best option did not receive the top reward (analogous to noise) was 0.2. Again, here adaptive learning was found in a regime of low volatility and intermediate noise, compatible with our theory (Fig. 7A).

The psychophysical experiment we performed further validated several prominent aspects of our theory that are common to both estimation and prediction tasks. We hope that future experiments will test several additional predictions. First, do people solve estimation and prediction problems according to the differences prescribed by our theory? For estimation problems, we showed that memory is not necessary when noise is low, regardless of volatility, whereas for prediction problems, memory is not required when both noise and volatility are low (as volatility increases, current evidence carries increasingly little information about the future, and thus it becomes useful to retain a long-term memory of the average source position). Moreover, complex strategies are useful over a wider region of the volatility/noise landscape for estimation problems than for prediction problems. Second, when and how do people use different forms of working memory (e.g., exponential or flat weighting)? Our theory predicts that, when volatility is not too high, flat weighting over past evidence is more useful at higher than lower levels of noise, and exponential weighting is most efficient at moderate noise, especially for prediction problems. Third, do subjects learn from recent evidence in conditions of high volatility and variable noise? Our theory predicts a transition between a domain where only prior information about the average source position is useful and a domain where that prior knowledge should be updated based on new evidence (Fig. 5). These two domains are separated by a roughly power-law curve in the volatility-noise plane, so that decreasing volatility increases the noise level beyond which learning from new evidence is useless. This transition curve is found for both estimation and prediction tasks. However, for prediction tasks the transition happens at lower noise levels because, when volatility is high, ongoing evidence is much less useful for predicting the future than for estimating the current source. Answering these questions will help to establish if and how trade-offs between accuracy and complexity govern the cognitive operations used to perform inference in the brain.

## Methods

### Gaussian change-point tasks

Models were tested using a Gaussian change-point task (Fig. 2) (Wilson et al., 2013; Nassar et al., 2010). Observations *x*_*t*_ were Gaussian distributed (*p*(*x*_*t*_) = *𝒩* (*x*_*t*_|*μ*_*t*_, *σ*^2^)) around a source located at an unknown mean position *μ*_*t*_. The mean position changed at random times, with probability *h* (the volatility parameter). At these change-points, the source was resampled from another Gaussian distribution 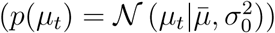. The goal of an observer was to infer the current position of the source *μ*_*t*_ from the history of observations up to time *t* (estimation problem), or to predict the position of the source at the next time step *μ*_*t*+1_ (prediction problem). The parameters, 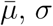, and *σ*_0_, were held constant in blocks and were assumed to be known to the observer; i.e., acquired after a sufficiently long exposure to the same environment. The ratio (*R* = *σ/σ*_0_) is the noise parameter of the process 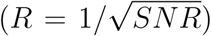.The volatility and noise parameters determined the statistical difficulty of the inference problem.

### Exact Bayesian inference

Here we derive expressions for 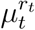 and *p*(*r*_*t*_|*x*_1:*t*_) to obtain the optimal Bayesian estimate of the current source position and the optimal Bayesian prediction of the next source position (text around eq. 2) (Adams & MacKay, 2007).

For Gaussian processes, the posterior probability of the source *μ*_*t*_ given run-length *r*_*t*_ is

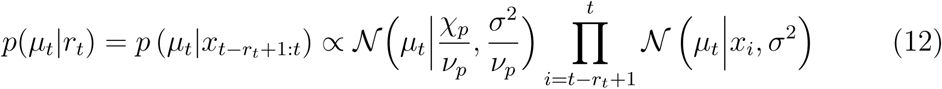

where we have used the Bayes rule 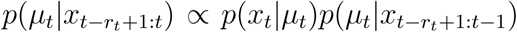 recursively. Note that 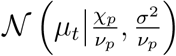 is the Gaussian prior distribution over *μ*_*t*_ with mean 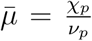 and variance 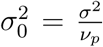. Using the relation 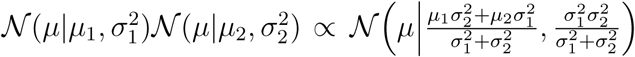 we obtain:

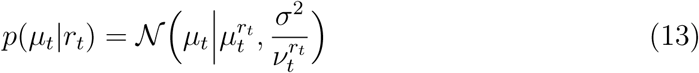

with

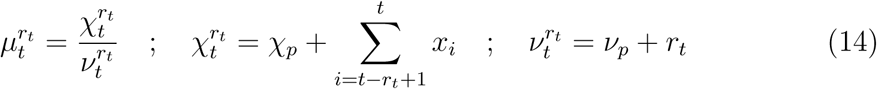

As expected for a Gaussian prior and a Gaussian likelihood, the posterior distribution (eq. 13) is also Gaussian.

The posterior probability of the run-length *r*_*t*_ given observations *x*_1:*t*_ can be computed recursively:

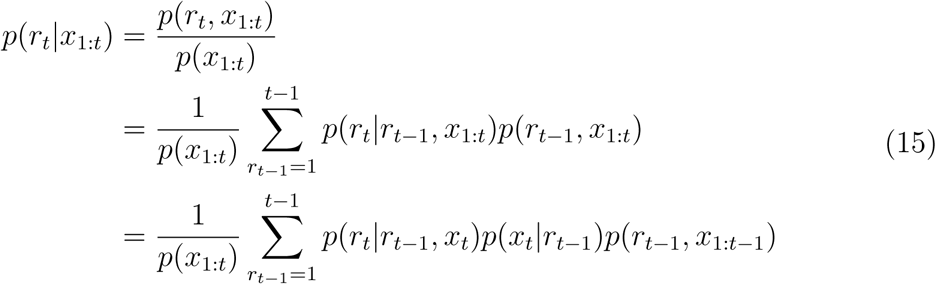

Because *r*_*t*_ = 1 if there is a change-point (“cp” below) at time *t, r*_*t*_ = *r*_*t*−1_ + 1 if there is no change-point, and change-points occur with constant probability *h*, we can rewrite *p*(*r*_*t*_|*r*_*t*−1_, *x*_*t*_) as:

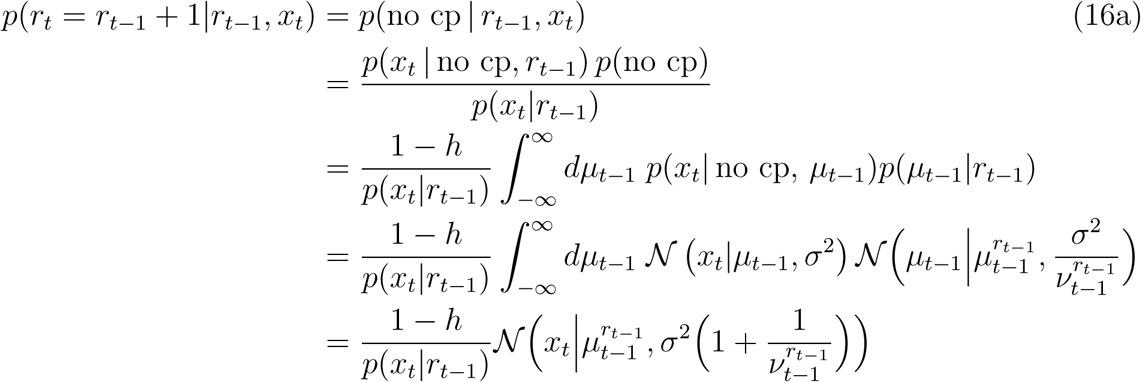

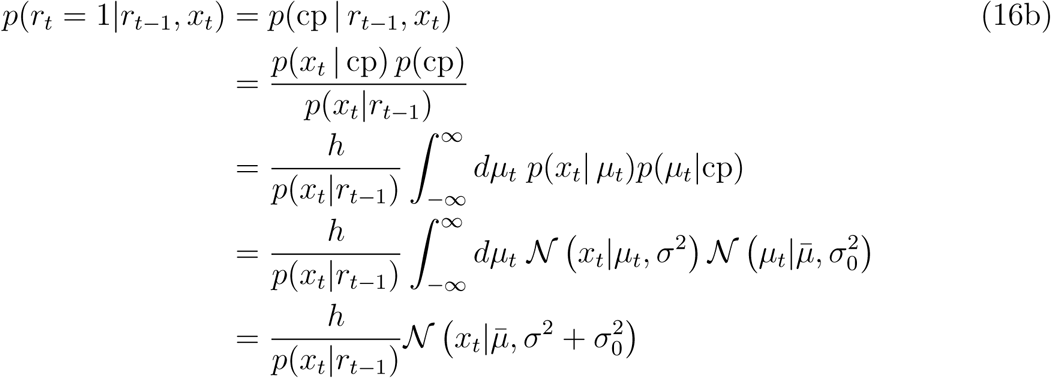

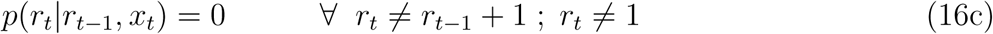

Substituting eqs. 16 into eq. 15 we obtain:

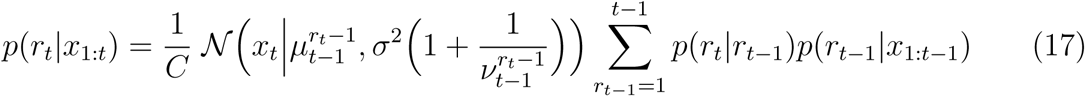

with *C* being a normalization constant, 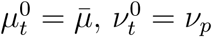 (for any *t*) and

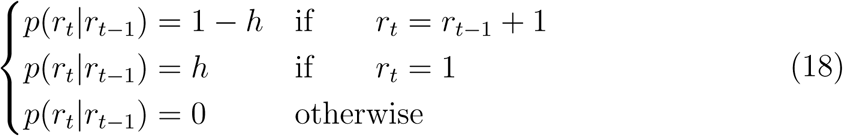

Eq. 17 simplifies to:

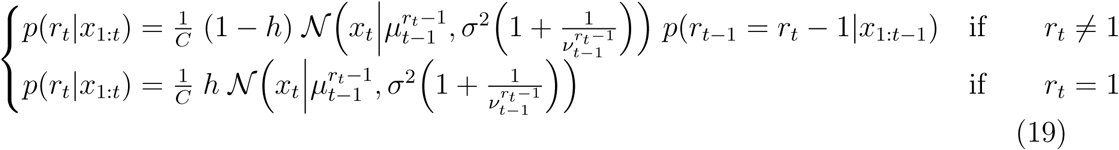

In conclusion, we can compute:

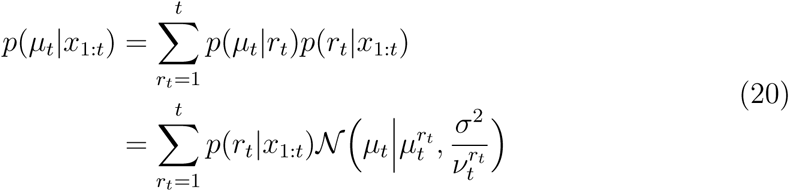

and the optimal (mean-squared-error minimizing) estimate of the source *μ*_*t*_ given the history of observations *x*_1:*t*_ is

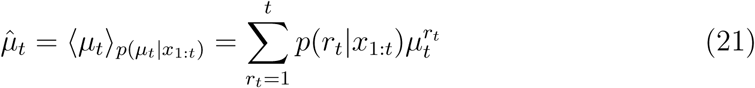

From *p*(*μ*_*t*_|*x*_1:*t*_) it is straightforward to derive the posterior probability distribution for the position of the source at the next time step:

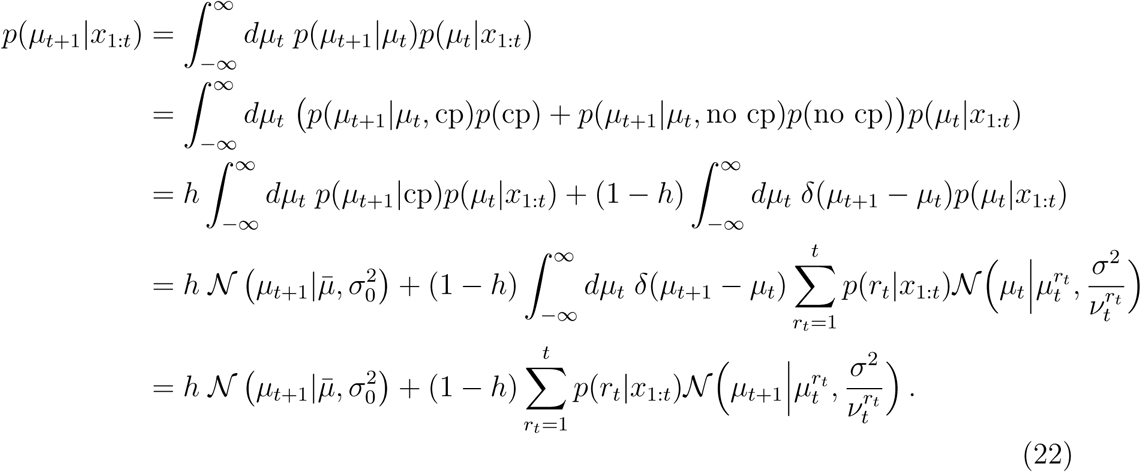

It follows that the optimal Bayesian prediction of *μ*_*t*+1_ given the history of observations up to time *t* is

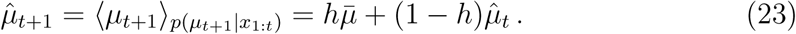

### Posterior probabilities in the Mixture Models

In the Mixture Models, the posterior probabilities of the run-lengths {*r*_*i*_}, *i* = 1,…,*N* are obtained as an approximation of the Bayesian posterior *p*(*r*_*t*_|*x*_1:*t*_) (compare with eq. 17 above)

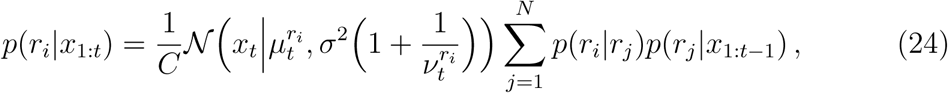

where the transition probabilities *p*(*r*_*i*_|*r*_*j*_) approximate *p*(*r*_*t*_|*r*_*t*−1_) of the exact Bayesian Model (Wilson et al., 2013, 2018):

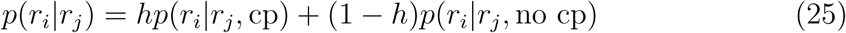

We sort the *N* model run-lengths in ascending order: *r*_1_ < *r*_2_ < *···* < *r*_*N*_. When there is a change-point, the Bayesian run-length drops to 1. This condition is approximated by resetting the model run-length to the smallest possible value *r*_1_:

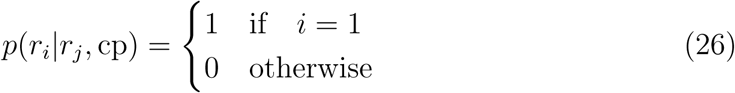

When there is not a change-point, the Bayesian run-length increases by 1. Given the finite number of run-lengths in the Mixture Models, the distance between any *r*_*j*_ and *r*_*j*+1_ is in general different from 1. To approximate the Bayesian transition, two cases are considered: (1) when *r*_*j*+1_ ≥ *r*_*j*_ + 1, the model run-length increases from *r*_*j*_ to *r*_*j*+1_ with a probability inversely proportional to the distance *r*_*j*+1_ − *r*_*j*_ and it remains constant with the complementary probability, so that the increase in model run-length is equal to 1 on average; (2) when *r*_*j*+1_ < *r*_*j*_ + 1, transition always occurs. More formally:

For all *j* < *N* :

If *r*_*j*+1_ ≥ *r*_*j*_ +1 then:

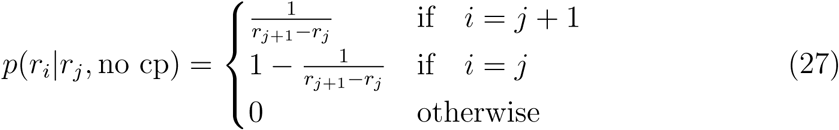

Else if *r*_*j*+1_ < *r*_*j*_ +1 then:

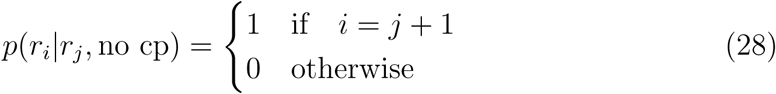

For *j* = *N* :

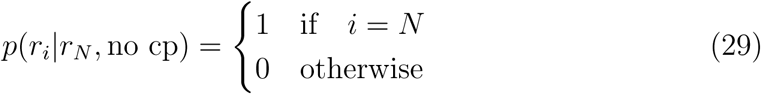

We have used Mixture Models with *N* = 2 units.

### Integration kernels and parameter reductions

The different models compute estimates 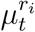 by using different integration kernels over past and present observations. The models based on Sliding Windows (both the *N* ≥ 2 adaptive and the *N* = 1 non-adaptive versions) compute 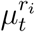 as in the Bayesian model (eq. 14) with *r*_*t*_ = *r*_*i*_. Eq. 14 can also be expressed as:

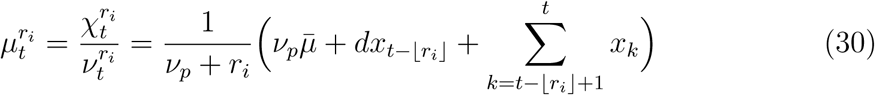

where we will think of the model run-length *r*_*i*_ as being allowed to take non-integer values in the mathematical expression to allow greater flexibility, and *d* = *r*_*i*_ − ⌊*r*_*i*_⌋ is the decimal part of *r*_*i*_. Eq. 30 corresponds to a sliding-window integration kernel over the most recent *r*_*i*_ observations, combined with the prior mean 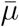. The relative weight of the prior mean with respect to each observation is 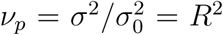: the larger the noise, the more the model relies on the prior mean as opposed to the empirical mean computed from the observations.

Eq. 30 can also be evaluated recursively as:

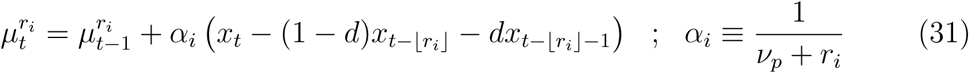

with initial condition 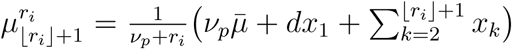. Note that the model still needs a memory that extends up to *r*_*i*_ time steps in the past. This model has an effective learning rate *α*_*i*_.

This working-memory load is reduced substantially in the Delta-Rule Models:

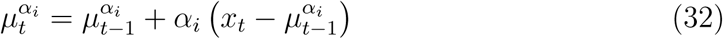

with initial condition 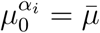 and learning rate *α*_*i*_ in the range [0, 1]. The delta-rule units are approximations of the sliding-window units in which the weighted average of the two observations occurring ∼ *r*_*i*_ time steps back in the past 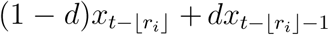 (eq. 31) is replaced by the unit estimate 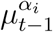 of the mean at time *t* − 1. This approximation reduces the working-memory demand to the previous time step only, at the cost of deteriorating the estimate of the source, especially at high noise or high volatility.

The integration kernel implemented by a Delta-Rule is an exponentially decaying kernel with time constant *τ* = −1*/* ln (1 − *α*_*i*_) (∼ 1*/α*_*i*_ for *α*_*i*_ ≪ 1):

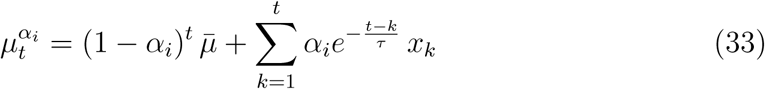

The Memoryless model further removes the dependence on the previous time step by estimating the source *μ*_*t*_ as a weighted average between the prior mean 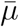 and the present observation *x*_*t*_ (Dirac-delta kernel):

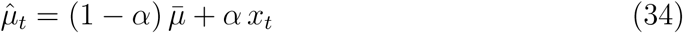

The parameters *r*_*i*_ and *α*_*i*_ of the models based on Sliding Windows and Delta Rules are optimized to minimize mean-squared error of the model estimates (or predictions), and their values vary with the environment-dependent parameters *h* and *ν*_*p*_ = *R*^2^. The optimal weight *α* of the Memoryless model is 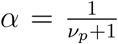 and is independent of volatility.

We observe a number of hierachical relationships between the models (Fig. 1 and Supplementary Fig. S1): first eq. 34 (with optimal *α*) coincides with eq. 30 under the constraint *r*_*i*_ = 1 (the Memoryless Model is nested in the Sliding-Window Model); furthermore, the simple Prior Model 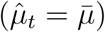 and the Evidence Model 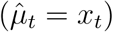 are obtained from both the Memoryless and the Delta-Rule Models by setting *α* = 0 and *α* = 1, respectively.

### Effective reduction to simpler nested models

We quantified the effective reduction of the adaptive Mixture Models into the associated non-adaptive nested models (single Sliding Window and single Delta Rule) in terms of a quantity that we called Alignment. The Alignment measures the angle between the eigenvector of the error Hessian matrix *H* (eq. 3) with the smallest eigenvalue (the “irrelevant eigenvector”), and the direction in parameter space between the optimal Mixture Model and the optimal non-adaptive nested model. Let us consider the two-parameter case. Let ***δα*** = ***α***^(1)^ − ***α***^(2)^, where 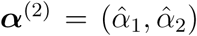 is the two-component vector of the optimal parameters of the Mixture Model, and 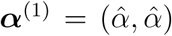 is the vector with both components equal to the optimal parameter of the non-adaptive nested model (Fig. 8). The vector ***δα*** is directed along the parameter transformation collapsing the best Mixture Model into the best nested Single-Unit Model. We then define

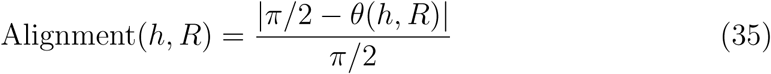

where 0 ≤ *θ* ≤ *π* is the angle between the irrelevant eigenvector and the direction of ***δα*** (Fig. 8). By definition, 0 ≤ Alignment ≤ 1 and is a function of volatility *h* and noise *R*. To reduce numerical noise, in Fig. 3, Redundancy and Alignment were averaged over ten instances of the Gaussian change-point process for each volatility and noise value.

**Figure 8:**
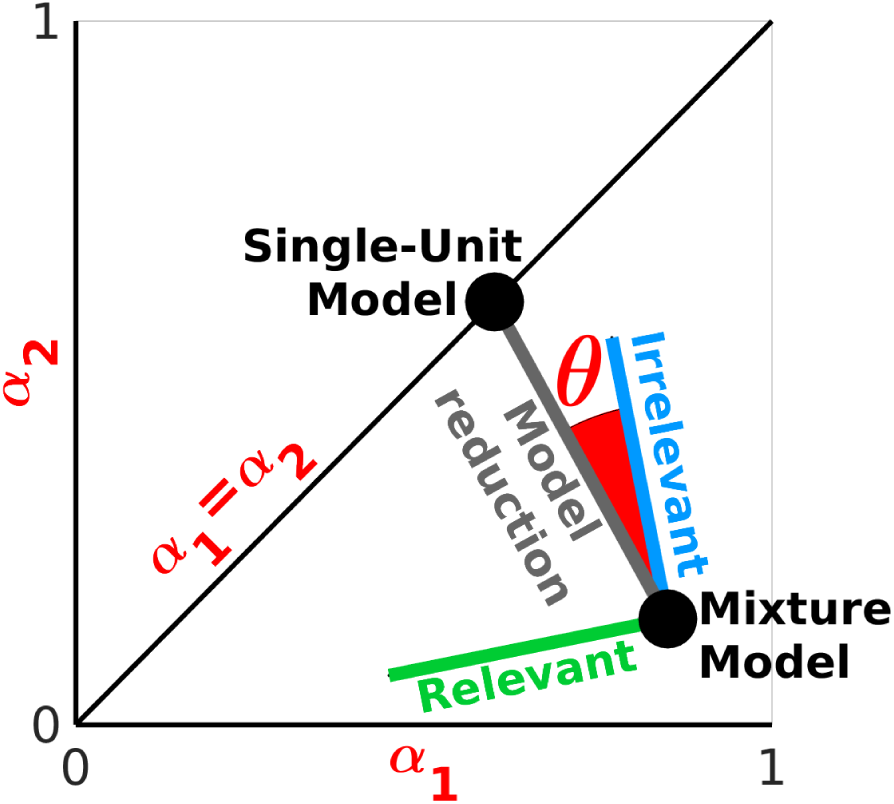
Computing Alignment. The axes represent the possible values of two learning rates, definining the parameter space for the Mixture Model; the diagonal *α*_1_ = *α*_2_ defines the parameter space for the Single-Unit Model. Black dots represent the optimal parameter configurations for the two models. Blue/green lines represent the eigenvectors of the Hessian (eq. 3) of the Mixture Model with the minimum/maximum eigenvalue, and constitute (under conditions of redundancy) the irrelevant/relevant degree of freedom of the model. The angle *θ* in eq. 35 is the angle between: (1) the irrelevant parameter direction of the Mixture Model; and (2) the direction of the parameter transformation ***δα*** (grey line) reducing the Mixture Model into its nested Single-Unit Model, in their optimal parameter configurations.

### Algorithmic complexity

The algorithmic complexity (eq. 5, Fig. 4A) is defined as the sum of a “reflexive” and a “reflective” cost.

The reflexive cost *C*_reflex_ of taking action once a decision has been made is not known, but because it is an equal constant for all models, its value does not influence the conclusions of this study (see Supplemental Information). We set *C*_reflex_ = 0.15 to obtain the power-law fits (Fig. 4), as this value optimizes the goodness-of-fit of the linear regression of log(ℐ) versus log(𝒞). Different values of *C*_reflex_ would just shift the power-law fits along the complexity axis by a constant.

The reflective cost is the sum of the computational and memory costs paid, on average, by each model to make an inference (estimation or prediction). We consider arithmetic operations (denoted by *A*), i.e., sums, subtractions, multiplications, divisions, exponentials, and square roots, and memory operations, i.e., writing (*W*), reading (*R*), and storing (*S*). For simplicity, we assign cost = 1 to each of these operations. Thus, the reflective cost reduces to the total mean number of operations per inference: ⟨*N*^*A*^⟩ + ⟨*N*^*W*^⟩ + ⟨*N*^*R*^⟩ + ⟨*N*^*S*^⟩, where we use the notation ⟨*N*^*i*^⟩ to indicate 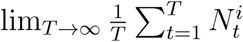, with 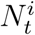 number of operations of type *i* ∈ {*A, W, R, S*} required in the *t*-th iteration (returning one inference) of the algorithm implemented by each model. More precisely, for memory operations, we define 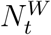 as the number of variables that have to be written into memory (at iteration *t*), 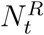 as the number of times each variable has to be read from memory (at *t*), summed over all variables, and 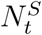 as the number of iterations (starting at *t*) during which each variable has to be kept in memory to make future inferences, summed over all stored variables.

Table 1 lists ⟨*N*^*A*^⟩, ⟨*N*^*W*^⟩, ⟨*N*^*R*^⟩, and ⟨*N*^*S*^⟩ for the estimation problem, for each of the seven models derived from the exact Bayesian strategy (Fig. 1). Below we explain how we determined these values, and how they can be readily converted into the corresponding values for the prediction problem. We will only indicate operations that are performed by the models in every inference, because 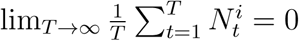 for one-off operations.

The Evidence Model returns each instantaneous piece of evidence 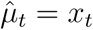, and does not require any computation or memory operation.

The Prior Model stores the prior mean 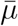 for one iteration at every *t* (⟨*N*^*S*^⟩ = 1) and reads it from memory (⟨*N*^*R*^⟩ = 1).

The Memoryless Model estimates the source position as 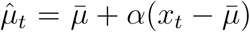, which requires ⟨*N*^*A*^⟩ = 3 (1 sum, 1 subtraction, 1 multiplication), ⟨*N*^*S*^ ⟩ = 2 (to store 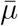 and *α*), and ⟨*N*^*R*^⟩= 3 (to read 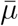, twice, and *α*, once). We stress that the name “Memoryless” is used to indicate that this model does not perform any integration of evidence over time, thus it does not require any memory of past observations or past inferences; however, the model maintains a memory of prior information.

The Delta Rule computes 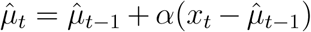, which involves the same number of algorithmic and memory operations as the Memoryless Model estimate, with the addition of one writing operation per iteration (⟨*N*^*W*^⟩ = 1), because the computation is recursive, requiring to write 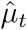 into memory at every *t* to compute 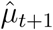. This one-time-step dependence allows the Delta Rule to integrate the evidence over time, unlike the Memoryless Model.

The Sliding Window computes 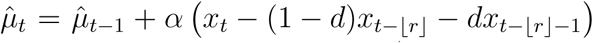 (with 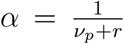), which requires ⟨*N*^*A*^⟩ = 7 arithmetic operations (1 sum, 3 differences, 3 products); ⟨*N*^*W*^ ⟩ = 2 operations to write, at every 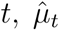 (necessary to compute the estimate at *t* + 1) and *x*_*t*_ (necessary to compute the estimates at *t* + ⌊*r*⌋ and *t* + ⌊*r*⌋ + 1); ⟨*N*^*R*^⟩ = 6 operations to read, at every 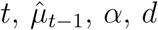 (twice), *x*_*t*−⌊*r*⌋_ and *x*_*t*−⌊*r*⌋−1_); finally ⟨*N*^*S*^⟩ = ⌊*r*⌋ + 4 operations to store, at every 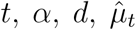 (for one iteration), and *x*_*t*_ (for a duration of ⌊*r*⌋ + 1 iterations). Because of the dependence on *r*(*h, R*) (the time scale of the sliding-window integration of past evidence that minimizes mean squared error), this model and its Mixture have complexity that depends on the environmental noise and volatility; for example, complexity increases with increasing noise to integrate observations over longer time scales, which allows more accurate estimates of the source. All the other models have complexity that is independent of noise and volatility, because they retain either no memory of past evidence (Evidence, Prior and Memoryless Models), or only a memory of the previous estimate (Delta Rule Models), regardless of environmental statistics.

In the Mixture models, each of the *N* units performs the same computations as the corresponding single-unit models. Thus, the contribution to the complexity of the Mixture models coming from the computations taking place in the single units reduces to the complexity of the single Delta Rule and single Sliding Window, respectively, when *N* = 1 (Table 1, first line of the respective slots). However, the largest contribution to the complexity of the Mixture models comes from the computations that combine the estimates provided by the *N* units into a single inference of the source (Table 1, second line of the respective slots, where the Heaviside function *H*_2_ = *H*[*N* − 2] vanishes for *N* = 1). These computations are necessary to obtain the adaptive probabilities *p*(*r*_*i*_|*x*_1:*t*_) of the *N* run-lengths at each iteration *t* of the algorithm, which are then used to weigh the estimates of the single units. In particular, for both Mixtures, the leading-order term of *N* ^*A*^ (7*N*^2^) comes from 2 summations, over *N* terms each, required to compute each of the *N* adaptive *p*(*r*_*i*_|*x*_1:*t*_) (eq. 24): (1) the summation over *N* run-lengths *j* appearing at the numerator of eq. 24 (which involves 6*N* − 1 algorithmic operations), and (2) the summation necessary to compute the normalization constant (which involves *N* − 1 algorithmic operations). The leading-order term of ⟨*N*^*R*^⟩ (6*N*^2^) comes from reading the terms in the same summations. ⟨*N*^*W*^ ⟩ scales as ∼ *N* (not as ∼ *N*^2^) because only the *N* probabilities *p*(*r*_*i*_|*x*_1:*t*_) are carried forward to the next iteration of the algorithm to compute the new *p*(*r*_*i*_|*x*_1:*t*+1_), whereas the individual addends of the summations mentioned above do not need to be memorized. Finally, the leading-order term of ⟨*N*^*S*^⟩ (2*N* ^2^) arises because computation of the adaptive *p*(*r*_*i*_|*x*_1:*t*_) requires maintaining in memory, at every iteration, the *N*x*N* matrices of the transition probabilities *p*(*r*_*i*_|*r*_*j*_, cp) and *p*(*r*_*i*_|*r*_*j*_, no cp) (eqs. 26 through 29).

The differences between the complexities of the Mixture of Delta Rules and the Mixture of Sliding Windows only involve terms of order 𝒪(*N*) and 𝒪(1), and come entirely from the computations taking place in the single units (note that the second line in the slots of Table 1 corresponding to the two Mixture models are identical).

The complexities in the prediction problem can be readily obtained from the complexities in the estimation problem, as follows. For the Evidence and Prior Models, predictions coincide with estimations, thus their complexity is the same as in Table 1. For all the other models, predictions are computed from estimations as 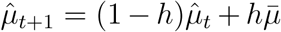. Thus, each prediction requires 4 more algorithmic operations than each estimation, 3 more reading operations (to retrieve from memory *h*, twice, and 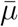, once), and either 2 more storing operations for the Delta Rule and Sliding Window (to store both *h* and 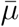), or just 1 more storing operation for the Memoryless Model (to store *h*, as this model already requires to store 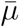 to obtain the estimate 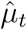) and for the Mixture Models (to store 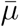, as *h* is already stored to estimate 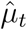).

### Psychophysics experiment

We recruited 169 subjects using the Amazon Mechanical Turk crowdsourcing website. The subjects performed an estimation task (which from pilot studies was less confusing than a prediction task) that used a generative process similar to the ones illustrated in Fig. 2. The task was presented as a card game. Before starting the game, subjects were given written instructions about the statistics of card and deck numbers. First, they were informed that deck numbers were picked in a 0-5000 range, that decks in the middle of the range (around 2500) were most likely, and decks at the extremes (around 0 and 5000) were least likely. Second, they were informed that each deck had cards with numbers that were near but not always the same as the deck number. Third, they were informed that (a) the randomness of the numbers in each deck, and (b) how often the decks switched without notice were both held constant within each block of trials but could change in the different blocks. The values of the randomness (noise) and the switching rate (volatility) were both explicitly indicated to the subjects via thermometer screen icons during task performance.

On each trial, the subject was shown a card number (corresponding to an observation *x*_*t*_ in Fig. 2) drawn from a card deck with Gaussian noise centered around the deck number. The deck number (corresponding to the mean *μ*_*t*_ in Fig. 2) was hidden to the subjects and changed at random times with a constant rate *h*. At the change-points, the deck number was resampled from a normal distribution with constant mean at 2500 (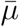 in Fig. 2). Non-integer values for card and deck numbers were approximated to the nearest integer. The ratio between the standard deviation of the card numbers around their deck number and the standard deviation of the deck numbers around their mean represents the noise parameter *R*. Subjects were asked to guess, on each trial, from which deck the card was being picked (i.e., to estimate the generative mean).

All 169 subjects were exposed to 40 training trials, in which we gave them the correct answer to familiarize them with the task. Training was followed by 360 test trials, in which we did not show the correct answer but provided feedback, after each trial, about the subject’s error relative to a weak and a strong competitor (the Evidence and the Bayesian models, respectively). Each subject performed 3 blocks of 400 trials in total (training + test). The blocks differed in terms of their values of noise or volatility (Fig. 7A): 85 subjects performed 3 blocks at constant volatility (*h* = 0.1) and variable noise (*R* = 0.01, *R* = 0.8, *R* = 4), and 84 subjects performed 3 blocks at constant noise (*R* = 1) and variable volatility (*h* = 0.08, *h* = 0.38, *h* = 0.8). Trials in which subjects input a number outside the 0-5000 range prescribed for the card decks, or responded in less than 100 ms, were excluded from further analyses.

### Data analysis

Values of adaptivity and working-memory load for subjects and models were obtained through the following analyses:

#### Integration kernels

For each subject and noise/volatility condition, we computed an integration kernel (linear weighting function) for each set of trials having the same lag from a change-point. Thus, we considered the set of subject responses 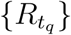 from all trials 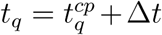, with Δ*t* a fixed lag, 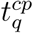 the *q*-th change-point trial, and *q* running from 1 to *M* (number of change-points in a given block occurring at 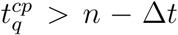, see below). From the response set 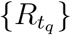 we estimated the subject integration kernel for the lag Δ*t*, by finding the weights *K*_*p*_, *K*_0_,…, *K*_*n*_ of the multiple linear regression model

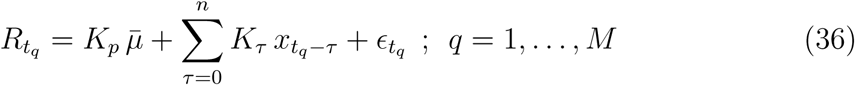

*K*_*p*_ (the weight given to the prior 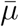) and *K*_0_,…, *K*_*n*_ (the weights given to the *n* +1 most recent observations) were obtained using the Matlab *lsqlin* function, which minimizes the sum of squared residuals 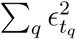 for the system of linear equations, with constraints 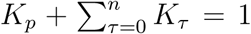 and 0 ≤ *K*_*i*_ ≤ 1, *i* = {*p*, 0,…, *n*}. We used *n* + 1 = 15 predictors (in addition to the prior) for the results in Fig. 7. Our conclusions were robust against changes in the number of predictors (Fig. S5). We estimated one integration kernel (set of weights) for each subject, block, and lag Δ*t*. We excluded only a few cases in which the linear regression model had fewer equations than predictors (because of an insufficient number of change-points at 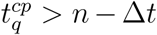), yielding underconstrained weights.

#### Integration time scales

For each integration kernel, we computed the normalized cumulative weight of the most recent *τ* observations

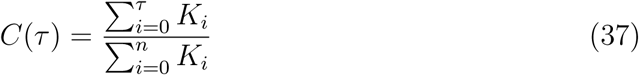

and the time scale at which this normalized cumulative weight reaches a fixed threshold *θ*

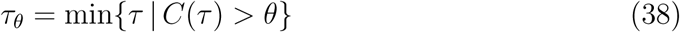

*τ*_*θ*_ represents an integration time scale. For the results of Fig. 7, we used *θ* = 0.8, so that *τ*_*θ*_ is the time scale that explains 80% of the subject’s integration over recently observed data. Conclusions were in general robust against changes in the threshold *θ* (Fig. S5). We computed one value of *τ*_*θ*_ for each integration kernel; i.e., for each subject, block, and lag Δ*t*. We estimated the standard error on each *τ*_*θ*_ by bootstrapping the regression model, eqs. 36. We used 200 bootstrap samples of the form 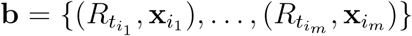, with 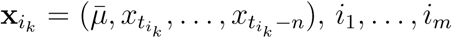 a random sample (with replacement) of the integers 1 through *M* (see Efron & Tibshirani (1994) for more details about the bootstrap procedure).

#### Adaptivity

For each subject and each block of trials, we computed adaptivity as the variance of *τ*_*θ*_ across all lags 0 ≤ Δ*t* ≤ *n* (because the integration time scale *τ*_*θ*_ can not be larger than the time scale *n* in the linear regression model). This metric of adaptivity quantifies how much the integration time scale changes as more and more card numbers are observed from the same card deck, in a given block of trials. To capture changes in individual adaptivity values across noise/volatility levels, we normalized each subject’s adaptivity in any given condition by the maximum adaptivity for the same subject across the three conditions, then we averaged the normalized values across subjects. The few subjects (3 for the variable noise conditions, 1 for the variable volatility conditions) for whom adaptivity was zero in all the three conditions were excluded from the analyses, because the normalized adaptivity was undefined. Standard errors on the average normalized adaptivity were computed as SEM 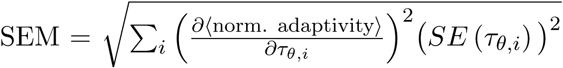 with the sum running over all the *τ*_*θ*_ obtained for different subjects and lags Δ*t*.

The theoretical estimates of adaptivity were obtained by considering the most-efficient model in each tested condition (Fig. 7A) and using the model simulated outputs, under the same sequences of observations shown to the subjects, in place of 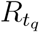 in eq. 36; *τ*_*θ*_ and normalized-adaptivity values were then obtained in the same way as the subject’s values. For the main bars in Fig. 7B, the most-efficient model was defined as the simplest model in our hierarchy with ℐ < 0.1 (as in Fig. 5B); theoretical predictions were qualitatively conserved across a relatively wide range of tolerance levels (e.g., between 0.02 and 0.2, see Fig. 7B, dashed gray lines).

#### Working-memory load

For each subject or most-efficient model and each block of trials, we computed working-memory load as the maximum *τ*_*θ*_ across all possible lags 0 ≤Δ*t* ≤ *n*; i.e., the maximum integration time scale (relative to the threshold *θ*) used for any given condition of noise and volatility. We then normalized this value by the maximum working-memory load across conditions for each subject and averaged the normalized values across subjects. Error bars were computed as for adaptivity.

## Acknowledgments

We thank Alex Filipowicz, Kamesh Krishnamurthy and Eugenio Piasini for many interesting discussions. We also thank Andrea Cavagna and Alessandro Ingrosso for pointing out a possible connection between one of our results and spin-glass systems. GT is supported by the Swartz Foundation and the Computational Neuroscience Initiative of the University of Pennsylvania. VB and JG are supported in part by NIH BRAIN Initiative grant R01EB026945. JG is also supported by R01 MH115557 and NSF-NCS 1533623.

## Author Contributions

G.T., V.B., J.G. developed the ideas and wrote the paper; all the authors designed the psychophysics experiment; T.D., C.P. performed the experiment; G.T. developed the theory and analysed the data.

