## Supplemental Information for "The complexity dividend: when sophisticated inference matters"

*Non-normalized versus normalized inaccuracy*

We define the non-normalized inaccuracy (Supplemental Figs. S3 and S4) of a model compared to the Bayesian ideal observer as

$$\mathcal{I}(h, R) = (E(h, R) - E_{Bayes}(h, R))/\sigma_0^2 \tag{1}$$

Both the normalized inaccuracy (eq. 6 in the main text) and the non-normalized inaccuracy (eq. 1) are invariant with respect to transformations that rescale the variance of the observations $\sigma^2$ and the variance of the source $\sigma_0^2$ by the same factor, so that the signal to noise ratio $1/R^2$ remains unchanged. Both definitions can be used, but emphasize different aspects of prediction and estimation accuracy. The normalization in eq. 6 of the main text defines inaccuracy as a relative quantity and thus emphasizes very accurate predictions when the ideal observer is very precise (small $E_{Bayes}$). By contrast, the quantity in eq. 1 treats inaccuracy as an absolute quantity, valuing accuracy in the same way regardless of how precise the ideal observer is.

*Optimal reflective cost is independent of reflexive cost*

When the reflexive cost $C_{\text{reflex}}$ changes, the optimal complexity shifts by the same amount:

$$\mathcal{C}_{opt}(h, R) = s + \left( \frac{a(h, R)\sqrt{b(h, R)}}{\sigma_r} \right)^{1/b(h,R)} \tag{2}$$

with $s = C_{\text{reflex}} - C_{\text{ref}}$, representing a shift from the reference value $C_{\text{ref}} = 0.15$ used to fit the model data. Therefore, the remaining reflective effort to maximize performance per unit complexity is unaffected by the reflexive cost, and the optimal strategy does not change.

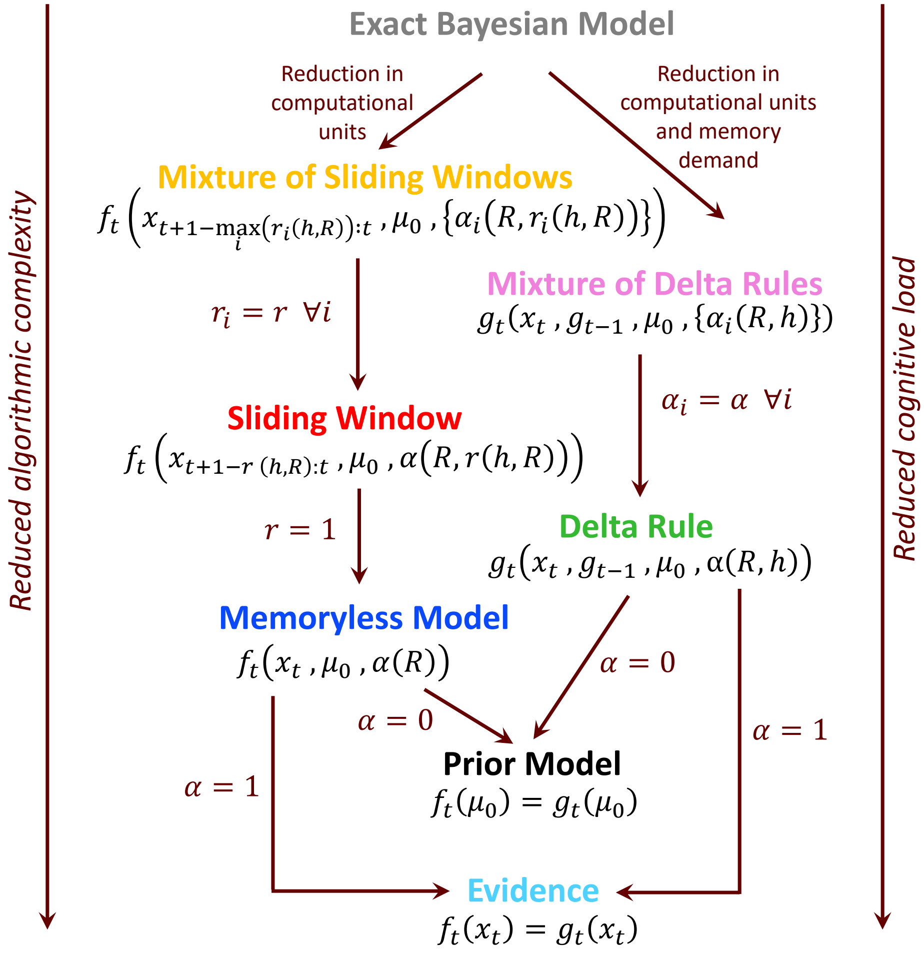


Figure S1: **Hierarchy of models: mathematical definition. Related to Fig. 1**. From the exact Bayesian strategy, two families of nested models can be derived, which compute the estimate $\hat{\mu}_t$ (formulas in black) through functions $f_t$ and $g_t$, respectively, of: (1) the observations ($x_{1:t}$); (2) the fixed parameters of the environment ($\bar{\mu}$, $h$, and $R$); and (3) the model-dependent "meta-parameters" (the run-lengths and the learning rates), which are themselves functions of the environment parameters. The transformations (arrows) that give rise to the two model families consist in progressive reductions of the meta-parameters (brown equations), which decrease computational complexity and cognitive load.

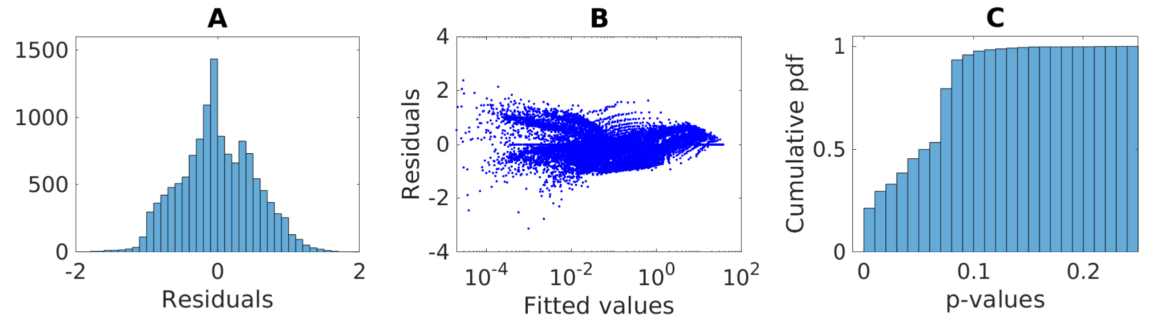


Figure S2: **Goodness-of-fit summary statistics for the power-law fits (linear fits in log-log scale) of inaccuracy $\mathcal{I}$ versus complexity $\mathcal{C}$. Related to Fig. 4**. The fits were obtained for a wide range of volatility and noise conditions ($0.02 \lesssim h \lesssim 0.85$ ; $0.1 \lesssim R \lesssim 6$). The distribution of residuals (**A**) is approximately normally distributed, and (**B**) does not show any specific dependence on the fitted values, indicating overall goodness of the fits. (**C**) The cumulative distribution of p-values for the F-test on the linear regression (log-log scale) for conditions in which more than two models have $\mathcal{I} \neq 0$ (finite F values) shows significance of the linear regression at the 10% significance level (or smaller) in over 95% of conditions.

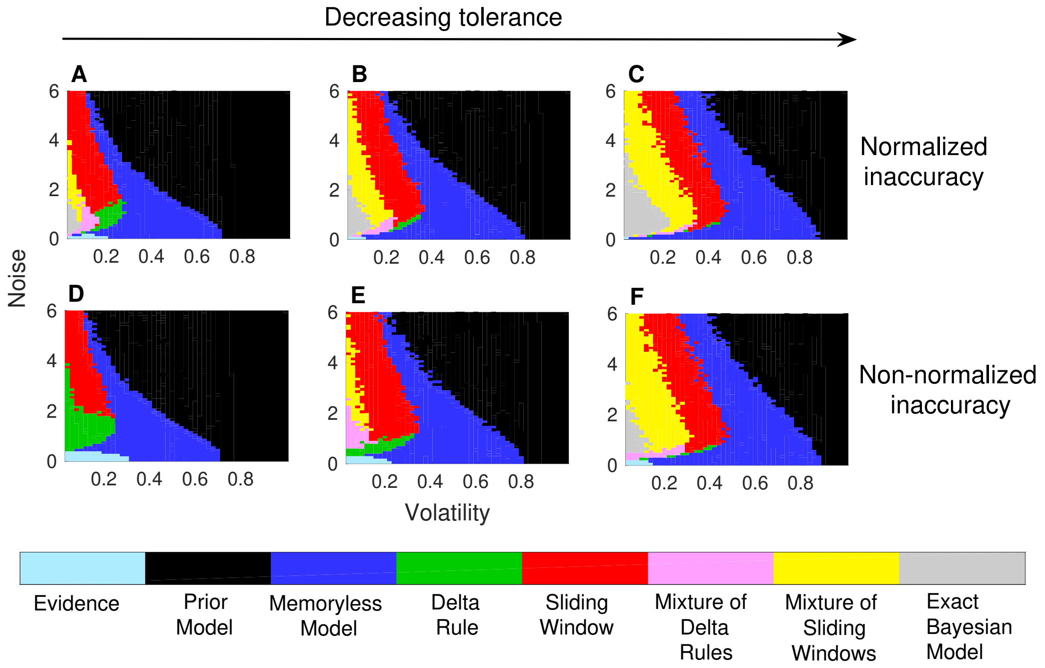


Figure S3: **The inverse-U relationship between the need for complex strategies and noise, at low volatility, is robust across criteria for assessing model performance and tolerance thresholds in the prediction problem. Related to Fig. 5**. The maps show, for each given volatility and noise combination, the simplest prediction strategy with $\mathcal{I}$ < tolerance. **A, B, C**: $\mathcal{I}$ = normalized inaccuracy (eq. 6 in the main text). **A**: tolerance = 0.1 (as in the main text); **B**: tolerance = 0.05; **C**: tolerance = 0.02. **D, E, F**: $\mathcal{I}$ = non-normalized inaccuracy (eq. 1). For comparison with the normalized inaccuracy, tolerance values in panels A, B, and C are mapped to $\langle E_{Bayes}\rangle_{h,R} \times$ tolerance$/\sigma_0^2$, yielding: **D**: tolerance = 0.089; **E**: tolerance = 0.044; **F**: tolerance = 0.018.

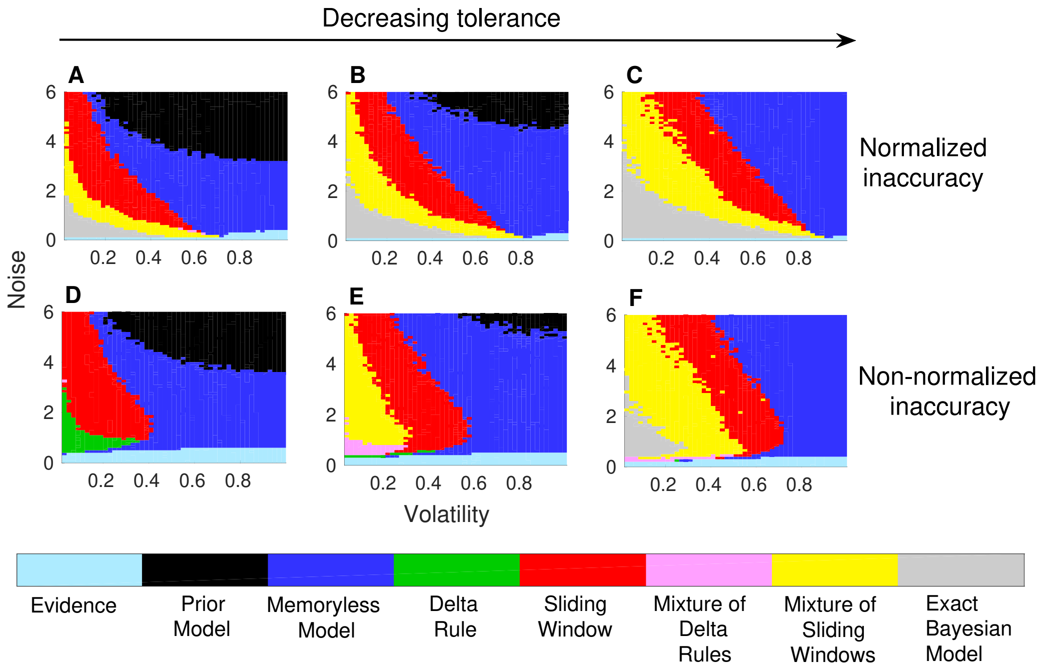


Figure S4: **The inverse-U relationship between the need for complex strategies and noise, at low volatility, is robust across criteria for assessing model performance and tolerance thresholds in the estimation problem. Related to Fig. 5**. Same definitions of inaccuracy and same tolerance thresholds as in Fig. S3. The inverted-U behavior of optimal complexity as a function of noise (see main text) is highly skewed at low noise when performance is measured in terms of normalized inaccuracy, whereas it is smoother (similar to the prediction case) when the inaccuracy is not normalized by the Bayesian model estimation error. This difference arises because the Baysian model is highly accurate at very low noise (yielding a skewed inverse-U structure when $\mathcal{I}$ is normalized) and absolute differences in errors between the Bayesian Model and simpler strategies are small in this limit (yielding a smooth inverse-U structure when $\mathcal{I}$ is not normalized).

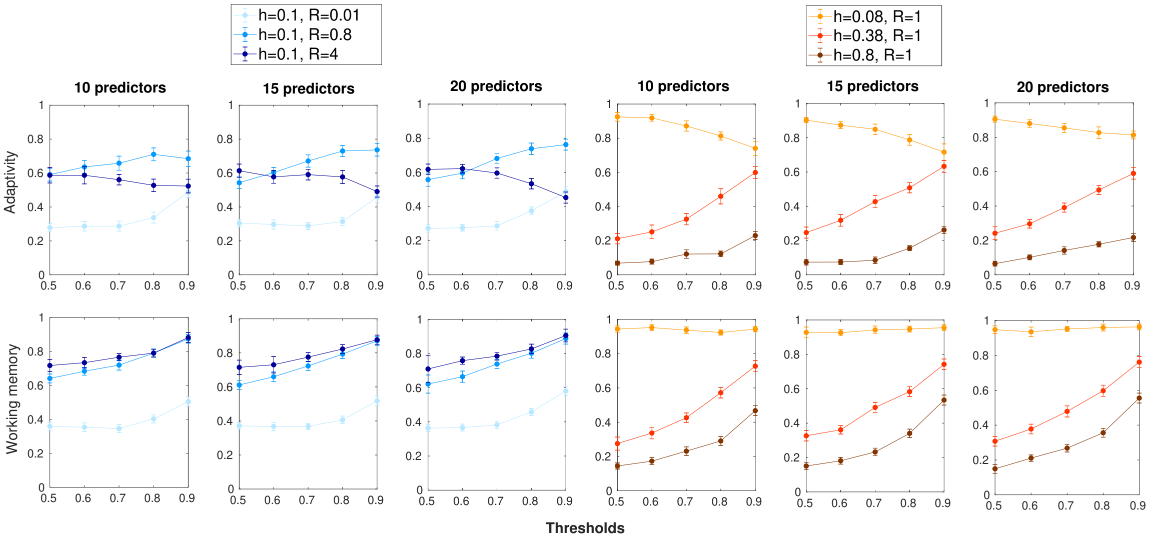


Figure S5: **Modulation of adaptivity and working-memory load with noise and volatility for human subjects is robust against changes in the number of predictors and thresholds $\theta$. Related to Fig. 7**. Mean normalized adaptivity and working-memory load ($\pm$ SEM) were computed across subjects, for different numbers of predictors in the linear regression model used to extract the subject integration kernels, eqs. 36 (Methods), and for different thresholds $\theta$ used to define the integration time scales $\tau_\theta$, eq. 38 (Methods). Changing the number of predictors and the threshold does not change, in general, the relative relationships predicted by the theory between adaptivity and working-memory values in the different noise and volatility conditions.
